# Single-cell FM streaming using genetically encoded protein oscillators

**DOI:** 10.1101/2025.02.28.640587

**Authors:** Rohith Rajasekaran, Thomas M. Galateo, Jing Pang, Zhejing Xu, Dennis T. Bolshakov, Elliott W. Z. Weix, Scott M. Coyle

## Abstract

Wireless devices use frequency modulation (FM) to reliably transmit information. Here, we establish a biochemical analogue of this paradigm using genetically encoded protein oscillators (GEOs) as carrier signals for real-time streaming of single-cell data. The GEO platform leverages evolutionarily diverse MinDE-family ATPase and Activator modules to generate fast synthetic protein oscillations in cells, where the waveform is controlled by both circuit biochemistry and intrinsic cellular physiology. This allows for diverse data encoding strategies, including noise-resistant FM reporters that broadcast proteasomal degradation or transcriptional dynamics; and autoencoder circuits that directly couple shifts in cell physiology to GEO waveform. Deploying GEOs in human embryonic stem cells allowed us to track single-cell developmental states in living populations during differentiation, revealing distinct patterns of spatiotemporal heterogeneity along endoderm, mesoderm, and ectoderm trajectories. The GEO platform establishes a dynamically controllable biochemical carrier signal, unlocking new high-fidelity FM data-encoding paradigms for continuous single-cell analysis.

## Introduction

Cells dynamically regulate gene expression, protein localization, and signaling state across different timescales to perform essential biological functions^1–4^. While genomic, transcriptomic, and proteomic methods can provide snapshots of single cell states^5–8^, the ability to follow the trajectories of individual cells in real time is critical to understanding how dynamic cellular and organismal behaviors are encoded and function^1,9,10^. These single-cell dynamics are often tracked microscopically using fluorescent reporters whose intensity or localization provides a proxy for the data of interest^10–16^. While powerful, these tools present challenges for extended tracking of single-cell dynamics and data aggregation, as arbitrary signal intensity varies across instruments and is sensitive to photobleaching and noise^17^. Moreover, the signals generated by traditional fluorescence-based tools lack metadata for identifying the underlying cellular source of the signal, making separation of overlapping signals in dense cellular environments difficult.

Electrical engineers overcome similar challenges in wireless communication by coupling data of interest to changes in the frequency of an easily measured carrier oscillation^18–20^. Because data is encoded in the temporal structure of the oscillation, frequency modulation (FM) has the advantage that measurements are in non-arbitrary units (ΔHz) and robust against noise and fluctuating intensity, making them consistent across different contexts and instruments. In addition, by assigning different carrier frequencies to different senders, multiple independent streams of FM data can be transmitted in the same medium and reliably unmixed using a range of powerful signal processing techniques. Extending FM data-encoding capabilities to biological measurement would provide a powerful paradigm for measuring single-cell dynamics and defining their biological consequences. However, this requires a biochemical analogue to the electromagnetic waves that serve as signals in radios and other wireless devices, as well as a detailed understanding of how the waveform can be biologically controlled and manipulated.

Recently, we found that synthetic protein waves can be generated in human cells through expression of the orthogonal *E. coli* divisome positioning system MinDE^21^. Here, an ATPase (MinD) and its ATPase-activator (MinE) self-organize fast ATP-driven protein waves that coherently propagate on membranes to fill the cell interior (Fig. 1A and Movie S1)^22–25^. When fused to fluorescent proteins, the resulting MinDE oscillations elegantly barcode single-cell identity and morphology with a unique frequency that can be analyzed using signal processing tools such as Fast Fourier Transform (FFT) and Continuous Wavelet Transform (CWT)^21^.

**Figure 1.**
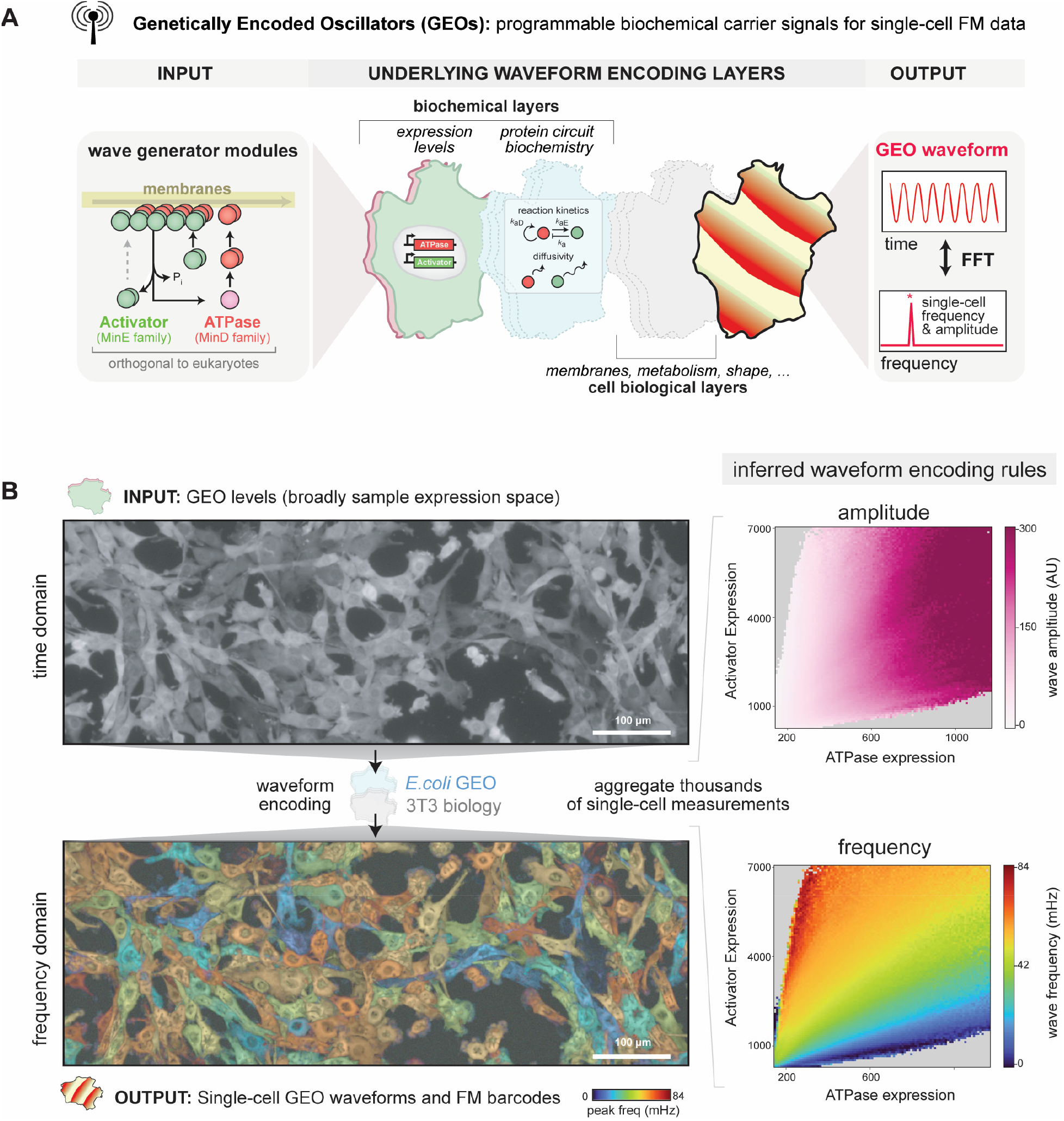
Genetically encoded oscillators (GEOs) generate biochemical carrier signals for single-cell FM streaming and data encoding. (A) Schematic of genetically encoded oscillators (GEOs) and their waveform encoding logic. GEOs are derived from MinD (ATPase) and MinE (Activator) bacterial divisome protein pairs that generate fast synthetic protein oscillations when expressed in human cells. These oscillations produce a characteristic single-cell frequency that can be analyzed using Fast Fourier Transform (FFT) and other signal processing tools. GEO waveform is encoded in both protein circuit biochemistry (expression levels and component sequence) and intrinsic cellular physiology, allowing for diverse data encoding strategies to be developed and deployed. (B) The underlying GEO waveform encoding rules in a specific experimental context can be inferred by measuring GEO frequency and amplitude across a wide range of ATPase and Activator expression levels. Aggregation of thousands of measurements across individual cells allows straightforward reconstruction of the underlying encoding logic in a phase portrait or 2D histogram that maps expression levels to waveform properties. See also Movie S1.

While these barcoded carrier signals provide each cell with its own unique “radio station”, the underlying frequency of this signal is fixed, leaving potential of these waves for streaming high-fidelity FM single-cell data untapped. Realizing this potential would require an expanded range of tunable frequencies and a clearer understanding of how expression levels, biochemistry, and cell physiology act as distinct layers that collectively encode a cell’s waveform. This would enable new paradigms for the design of FM streaming circuits that read out specific activities of interest or encode shifts in cell physiology directly in wave dynamics.

Here, we develop a modular platform for building and applying genetically encoded protein oscillators (GEOs) for real-time FM streaming of single-cell data. The GEO platform leverages evolutionarily diverse MinDE-family ATPase and Activator modules to expand the range and programmability of frequencies for data encoding. Through experiments, simulations, and modeling, we define how a cell’s waveform is controlled by expression levels, circuit biochemistry, and intrinsic cellular physiology. This allowed us to develop diverse single-cell streaming strategies, including noise-resistant GEO-FM reporters of proteasomal degradation or transcriptional dynamics; and GEO-autoencoder circuits that directly couple changes in cell physiology to GEO waveform. We demonstrate the utility of the GEOs in human embryonic stem cells (hESCs), tracking single-cell developmental states in living populations during differentiation. These FM-barcoded GEO signals could be projected directly onto microscopy images to reveal spatiotemporal state heterogeneity during differentiation into endoderm, mesoderm, and ectoderm lineages. The GEO platform establishes a dynamically controllable biochemical carrier signal, unlocking high-fidelity FM streaming paradigms for continuous single-cell analysis.

## Results

### Divergent GEO ATPase and Activator modules expand frequency bandwidth and enable predictable design of GEO waveform

The *E. coli* ATPase MinD and its activator MinE generate fast synthetic protein waves that propagate across membranes when expressed in human cells, providing a compact and orthogonal genetically encoded oscillator (GEO) that could serve as a carrier for FM transmission of biological data^21^. A cell’s observed waveform is encoded in the interplay between multiple different layers: the expression levels and biochemical properties of the ATPase and activator modules that produce the waves; and the cell’s internal physiology, which provides the membranes, structure, and metabolism that support wave propagation (Fig. 1A)^21–23^. While expression levels and GEO waveform are visible and can be measured directly, effects arising from different protein sequences or changes to cell physiology must be inferred. This can be achieved experimentally by constructing a phase portrait obtained from imaging thousands of cells, providing a statistically supported visual landscape for how ATPase and activator levels map to frequency and amplitude in a specific experimental context (Fig. 1B and Movie S1).

To expand GEO utility and programmability, we took an evolution-guided approach. We hypothesized that divergent MinDE-family ATPase/activator pairs across different bacterial species would possess altered reaction rates, stability, or membrane preferences that would change their waveform encoding rules^26^. Phylogenetic analysis of thousands of species-matched MinDE pairs queried the evolutionary diversification across ATPase and activator modules (Fig. 2A, Fig. S1). While the ATPase/activator structure and binding interface was generally conserved, we observed extensive sequence divergence within the individual proteins, especially in their membrane targeting sequences. Thus, ATPases and activators could likely act as separable modules for constructing GEOs with different quantitative behaviors and altered programmability.

**Figure 2.**
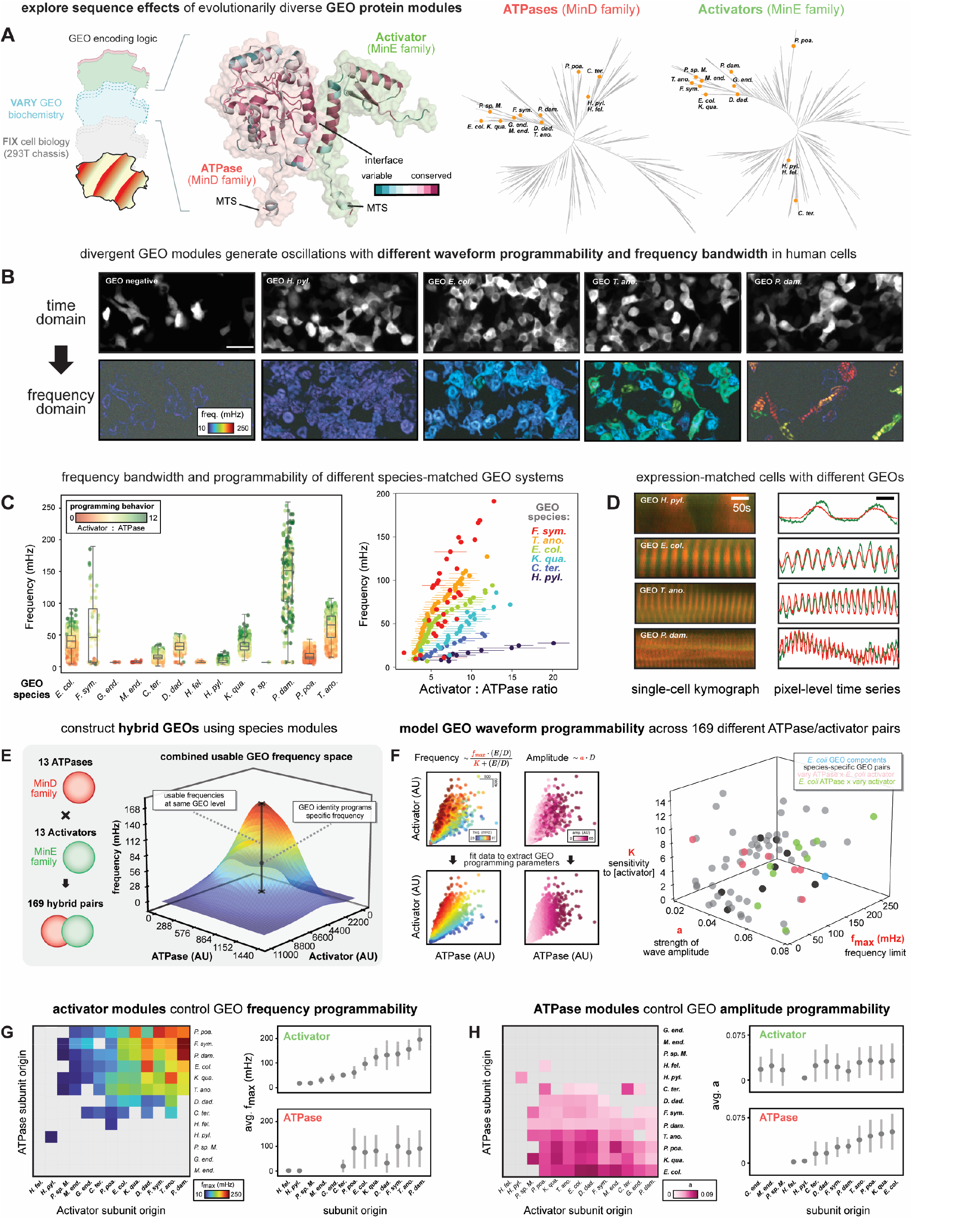
Divergent GEO ATPase and Activator modules expand oscillation bandwidth and enable predictable design of GEO waveform. (A) The effects of protein sequence and component biochemistry on GEO waveform encoding is explored by testing different evolutionarily diverged MinDE components in a common 293T background. ConSurf scores for GEO sequence conservation were derived from phylogenetic trees and multiple sequence alignments and mapped onto an AlphaFold2 structure prediction of thre *E. coli* GEO dimer. 4585 unique MinD ATPase sequences and 4331 unique MinE Activator sequences were aligned using ClustalW. Phylogenetic trees were built using the JTT matrix for maximum likelihood, and rapid bootstraps (100 replicates) with gamma substitution rates were used for tree construction. 13 selected species-matched GEO pairs are annotated on the resulting tree. See also Fig. S1. (B) Representative widefield fluorescence images and associated power spectrum images false-colored by frequency taken from 293T cells transiently transfected with different species-matched GEO pairs (GEO negative pair: mCherry and GFP expression). See also Fig. S2. (scale bar: 50µM). The representative variants in the panel are arranged to show a progression from slower (more blue) to faster (more red) frequency encoding behavior. (C) Left: swarm plot showing single-cell frequency quantification across a population of 293T cells expressing the respective GEO species-matched pair. Each point in the distribution corresponds to a cell, colored by its associated [Activator] to [ATPase] expression ratio. Right: scatter plot showing the relationship between [Activator] to [ATPase] expression ratio and GEO frequency for a subset of GEO species variants from the left panel. Points depict the expression ratio (mean value ± standard deviation) associated within each FFT-defined GEO frequency bin. See also Fig. S3. (D) Representative kymographs and pixel-level time series of GEO expression-level matched cells for different species-matched GEO pairs. (E) The ATPase and Activator components of a GEO pair act as separable modules for engineering oscillation waveform. 169 pairwise combinations of the 13 species-matched ATPase and activator pairs from (C) were tested. Individual frequency phase portraits associated with each pairwise combination were aggregated to generate a global accessible design space. A 3D surface visualizing the range of accessible frequencies at any GEO expression level using any GEO pairwise combination is shown. (F) GEO waveform frequency and amplitude encoding behavior can be phenomenologically modeled using simple mathematical relationships to extract intuitive programming parameters. A 3D scatterplot of *f*_*max*_, *K*, and *a* for 76 functional GEO systems is shown. See also Fig. S5-6. (G) Hierarchically clustered heatmap of the *f*_*max*_ or (H) *a* parameters from (F) for 169 pairwise GEO combinations. To identify trends, average parameter values were calculated for all ATPase pairings with a given activator (to test role of ATPase); or all Activator pairings with a given ATPase (to test role of Activator). For *f*_*max*_, the subunit origin axis is sorted by ascending average *f*_*max*_ of the Activator identity, while for *a*, the subunit origin axis is sorted by ascending average *a* of the ATPase identity.

To explore this, we selected 12 species-matched ATPase/activator pairs with varying levels of sequence similarity to *E. coli* MinDE for phase portrait analysis in 293T cells (Fig. 2B, Fig. S2-3). Of the 12 new GEO pairs tested, 9 robustly supported synthetic wave formation. Strikingly, many of these new GEOs exhibited altered waveform encoding behavior including shifts in the expression levels that supported oscillation and the range of frequencies that could be achieved (Fig. 2C, Fig. S3). For example, over the same range of [Activator] to [ATPase] ratios, the *F. symbiotica* GEO had wide bandwidth (11-169 mHz) compared to *E. coli* (25-58 mHz), while *H. pylori* GEO bandwidth was narrow (8-16 mHz). Expression-matched cells harboring different GEO pairs can thus encode oscillations with completely different frequencies (Fig. 2D).

Because the ATPase/activator interaction interface appeared conserved across these variants, we reasoned that hybrid GEO behaviors could likely be created by combining ATPase and activator modules from different species. We tested 169 pairwise combinations of the 13 ATPase and activator species, of which 76 hybrid GEOs produced oscillations in 293T cells that were robust enough for phase portrait analysis. Aggregating these data across all functioning GEO pairs provides a global design manual that can be computationally queried to identify modules that program a target waveform at a given expression level (Fig. 2E). This design space provides an accessible GEO bandwidth spanning 8-169 mHz at matched expression levels; and frequencies as high as 205 mHz using more specific configurations. To test the generality and transferability of the variant design space, we analyzed the behavior of the same panel of 169 pairs in a budding yeast chassis (Fig. S4). In this setting, we observed similar qualitative and quantitative trends to the 293T data, suggesting our portfolio of GEO ATPases and Activators has broad utility for waveform programming across different cell lines and diverse eukaryotes.

To understand how the different ATPase and Activator sequences contribute to GEO waveform, we fit a phenomenological model to each GEO’s phase portrait and used the fit parameters for quantitative comparison and trend analysis of module variants (Fig. 2F, detailed in Fig. S5). Oscillation *frequency* was treated as a hyperbolic response to the [Activator] to [ATPase] ratio, with *f*_*max*_ describing the maximum frequency achievable by a given GEO and *K* describing the sensitivity of GEO frequency to changes in [Activator]. Oscillation *amplitude* was modeled as a linear response to [ATPase] with slope parameter *a*. An associated wave energy can restrict fit predictions only to the regions of expression space where oscillations were experimentally observed to occur (Fig. S6). Using this approach, we reduced the description of each GEO’s programming rules and phase portrait down to three intuitive parameters.

A scatterplot of *f*_*max*_, *K*, and *a* for the 76 functional GEO systems revealed a wide range of waveform encoding behaviors (Fig. 2F). While species-matched ATPase/activator pairs expanded frequency programmability, inter-species hybrids accessed a significantly wider range and breadth of parameter space. Notably, only interspecies hybrids exhibited larger *K* values, allowing oscillations to be generated and tuned across a much wider range of expression levels. This feature is likely a consequence of imperfections at the binding interface between ATPase and activators that have not co-evolved.

Our hybrid screen analysis also revealed how the identity of ATPase and activator modules provides separable, independent control over specific features of the GEO waveform. Activator identity was the primary controller of a GEO’s frequency bandwidth *f*_*max*_ (Fig. 2G). For example, GEOs constructed using the *P. damselae* activator consistently oscillate faster than GEOs with the *E. coli* activator. Sorting activator modules by their average *f*_*max*_ across all hybrids provides a hierarchy of activator strengths for predictable design of faster or slower GEOs. In contrast, ATPase identity was the primary controller of a GEO’s amplitude parameter, providing predictable design of stronger or weaker oscillations (Fig. 2H).

Collectively, our ability to build many functional GEOs from a small set of GEO modules demonstrates the ease with which waveforms can be systematically programmed. These ATPase and activator modules provide a starting point for engineering more complex GEO circuit architectures for single-cell data-encoding applications.

### Multi-component GEOs provide compositional control of oscillation frequency

Because our variant screening found that different activator modules can cross-react with the same ATPase, we hypothesized that multi-component GEOs would also be functional, potentially providing an inroad for compositional control of GEO waveform and frequency (Fig. 3A). However, it is not obvious whether the biochemical mechanism underlying GEO wave propagation would support multiple competing components or how the resulting oscillations would behave.

**Figure 3.**
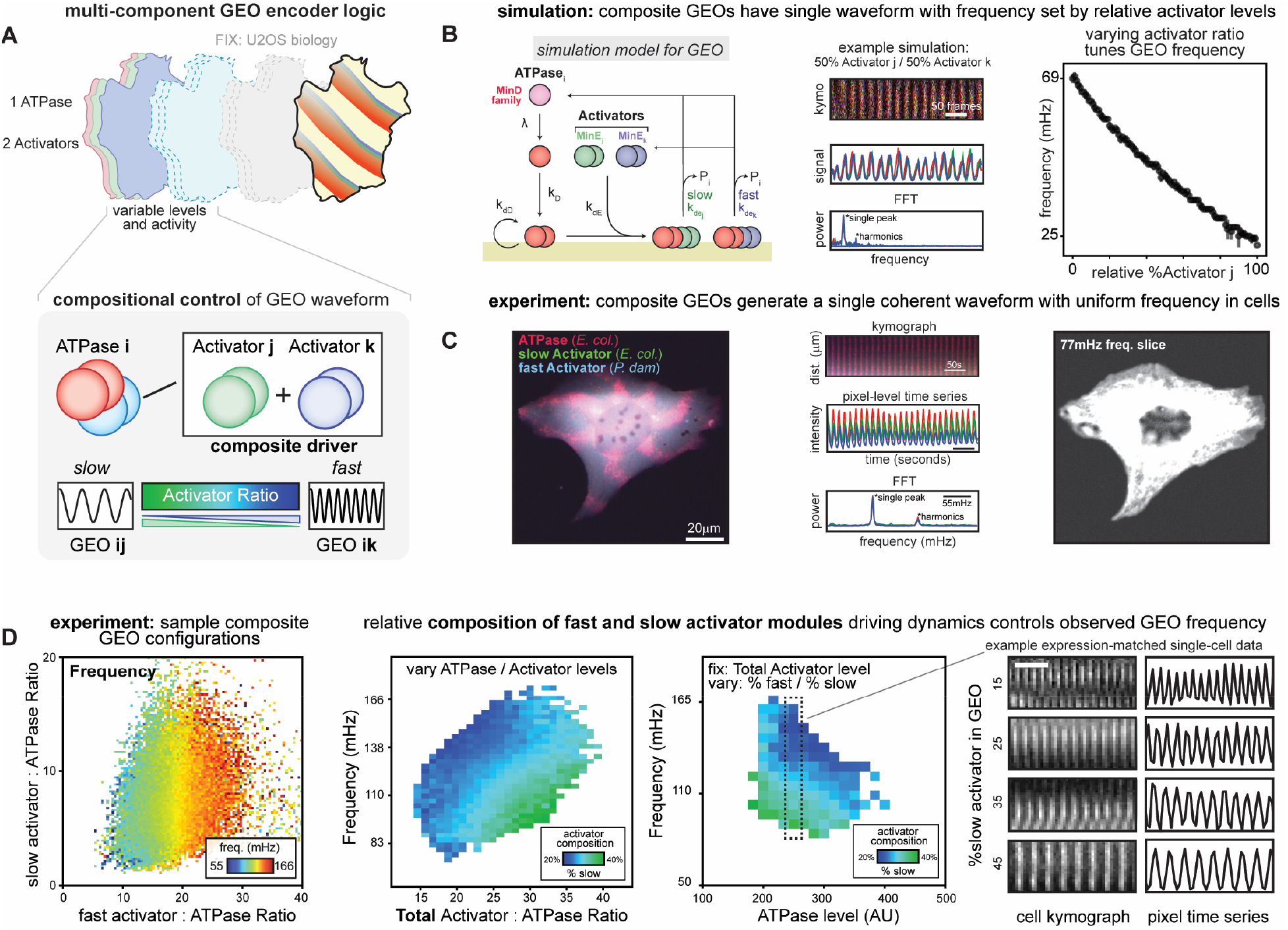
Multi-component GEOs provide compositional control of oscillation frequency. (A) Schematic and encoding logic for a multi-component GEO built using a single ATPase and a composite driver composed of a mixture of two competing Activators. (B) Simulation of multi-component GEO circuits. Left: diagram of the “skeleton model” of the multi-component GEO system used for simulation, depicting the biochemical species and the reactions that occur. The labels for the molecular species and rate constants associated with the formal description of the simulation equations are indicated. Center: a representative kymograph, pixel-level time series, and the associated FFT power spectrum for an example simulation of 50% Activator j and 50% Activator k is shown. Right: a set of simulations with varying % Activator j was performed. Peak frequency for each respective % Activator j and associated standard deviation is shown (10 replicates). See also Fig. S7. (C) Experimental validation of predicted multi-component GEO single waveform behavior. Representative fluorescence image of a U-2 OS cell transduced with a multi-component GEO system composed of an ATPase (*E. coli*), a slow (*E. coli*) activator, and a fast (*P. damselae*) activator. A representative kymograph, pixel-level time series, and the associated FFT power spectrum for this cell is shown. The cell appears on the 77 mHz frequency slice of the image power spectrum. See also Movie S2. (D) Aggregating single-cell multi-component GEO circuit behavior across thousands of U-2 OS cells reveals rules for compositional control of GEO waveform. A multi-dimensional phase portrait showing the observed relationship between relative protein expression levels and circuit frequency is shown. The [Total Activator] to [ATPase] expression ratio is a major determinant of GEO frequency. The frequency-scaling behavior for multi-component GEOs with different activator compositions is shown. The frequency dynamics for fixed [Total Activator] level and varying [ATPase] level are shown. Representative kymographs and pixel-level time series for [Activator] and [ATPase] expression-level matched cells with different compositions of Total Activator are shown. (scale bar: 50 seconds)

To gain intuition, we extended a stochastic model of GEO wave propagation in mammalian cells^21,27,28^ to include two chemically distinct activator modules: one that produces slow oscillations (Activator j); and one that produces fast oscillations (Activator k) (Fig. 3B, detailed in Fig. S7). Despite no explicit interaction between the two activators, simulations containing 50% Activator j and 50% Activator k produced a single composite waveform in which the two activators oscillated in phase. This coherence appears to arise through indirect coupling of activators via their shared ATPase node, as simulations using fully decoupled GEOs produced two separable frequencies (Fig. S7). To understand how the amounts of Activator j and Activator k control the waveform, we performed simulations that titrated the percentage of Activator k in the system from 0 to 100% at steps of 1% (Fig. 3B). We found that the wave frequency was smoothly titrated by the relative composition of the two activators. Thus, for this multi-component GEO architecture, the two activator species can be treated as a composite driver of oscillation frequency whose overall activity is controlled by relative levels of each activator.

We experimentally validated this multi-component GEO control logic in U2-OS cells, pairing slow (*E. coli)* and fast (*P. damselae*) activators with a single ATPase (*E. coli)*. Consistent with our modeling, the multi-component GEO produced a single waveform at one frequency with the two activator modules co-oscillating in phase (Fig. 3C). By sampling a wide range of expression levels across all three GEO components, we obtained a multi-dimensional phase portrait for how GEO composition programs waveform (Fig. 3D, Fig. S8). As in our modeling, the relative composition of the two activator species set the frequency of the oscillation, thus providing a potential strategy for building frequency-modulation GEO circuits.

### Engineering GEO-FM barcoded data streaming circuits for single-cell dynamics

The ability to tune the frequency of a GEO using compositional control suggests a strategy for building frequency-modulation (FM) barcoded streaming circuits (Fig. 4A). In these GEO-FM circuits, the constitutive expression of a “slow activator” (*carrier*) generates a constitutive GEO carrier oscillation barcode inside the cell. Data can be encoded in GEO frequency modulation by coupling the availability of a second “fast activator” (*encoder*) to any cellular activity of interest. Critically, these FM data streams would provide barcoded measurements of cellular state in non-arbitrary units (ΔmHz) that are transferable across instruments and insensitive to photobleaching and intensity fluctuations^18,19,29^.

**Figure 4.**
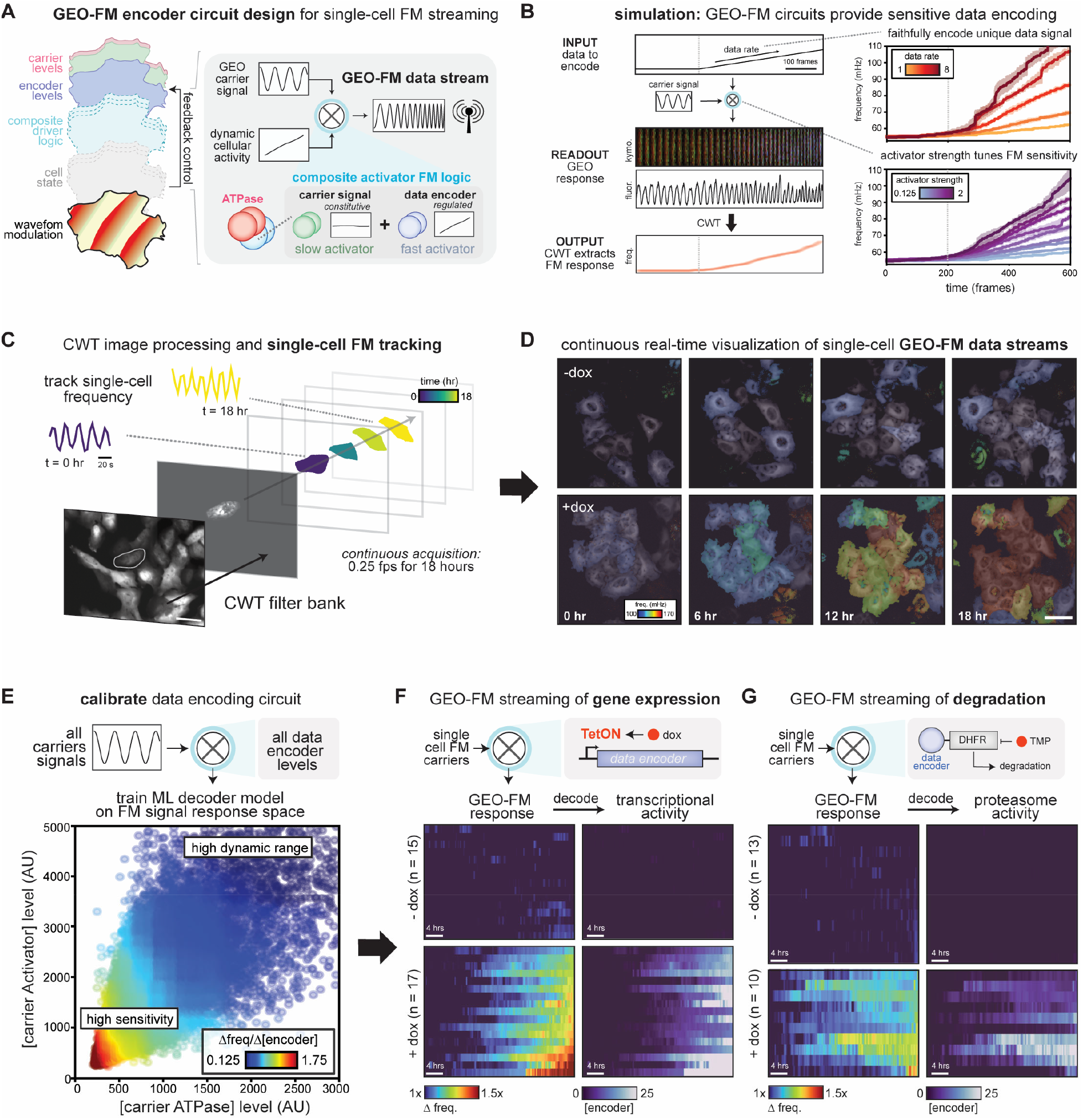
Engineering GEO-FM barcoded data streaming circuits for single-cell dynamics. (A) Schematic of GEO frequency modulation (FM) data encoder logic. A stable carrier signal is generated via constitutive expression of a slow GEO ATPase/Activator pair. Potent FM waveform modulation is achieved by coupling the expression or availability of a FAST Activator module to changes in cell state or physiology. (B) Simulated behavior of GEO-FM encoding circuits across different component parameterizations. The “skeleton model” from Fig. 3B was used to simulate and prototype GEO-FM circuit design. A simulated carrier waveform, composed of an ATPase and a carrier activator, was provided input data in the form of a ramped increase of an encoder Activator into the system. The simulated waveform was tracked to measure GEO response, and CWT was used to follow the instantaneous FM response over time. Simulations with different input data rates and different encoder activator strengths are shown. (C) Experimental workflow for collecting and analyzing single-cell GEO-FM data streams over time (detailed in Fig. S10 and Movie S3). Starting from a raw single-channel fluorescence microscopy time series, individual cells are segmented and tracked over time. The resulting masks are used to isolate and quantify each cell’s instantaneous frequency over time using a CWT filter-bank. Representative fluorescence and CWT-processed images, pixel-level oscillation dynamics, and GEO-FM frequency dynamics for a representative cell from the experiment in Fig. 4B processed using this workflow are shown. (D) Representative images visualizing the FM response of a population of U-2 OS cells bearing a doxycycline-inducible GEO-FM transcriptional encoder in the presence or absence of doxycycline. Data streams are visualized by overlaying the pixel-level instantaneous frequency estimate from a continuous wavelet transform (color coded by frequency) on to the associated pixel-level signal amplitude. Snapshots at different time points during an 18-hour time course are shown for the induced and uninduced population. See also Movie S3. (scale bar: 50µM) (E) Quantitative calibration of GEO-FM circuit responses. A machine-learning model was used to map changes in a GEO-FM carrier frequency to unknown data encoder levels. The model was trained using aggregated measurements from thousands of U-2 OS cells that sample a broad range of ATPase, carrier activator, encoder activator levels, and their corresponding oscillation frequency (detailed in Fig. S9). The resulting ML-predicted FM response sensitivity across different ATPase and carrier Activator expression levels is shown. (F) Engineered GEO-FM circuits can encode and stream changes in protein levels arising from dynamic transcriptional activity or (G) proteasomal degradation. To encode transcription activity, a genetic circuit placing an encoder Activator under the control of the doxycycline-induced TetON promoter was used. To encode proteasomal degradation dynamics, a protein circuit placing an encoder Activator under the control of a TMP-stabilized DHFR degron was used. Single-cell data streams were collected in the presence or absence of inducer at 0.25 FPS for 18 hours and analyzed using the analysis workflow from Fig. 4C. The frequency-barcoded FM responses for cells with different GEO-FM carrier signals are shown, ordered by weakest to strongest final FM response. These FM responses were converted to an associated change in the activity of interest (encoder levels) using the ML-model from Fig. 4D. See also Fig. S10 and Movie S3.

To build intuition for GEO-FM circuit design, we first modeled the behavior of different data encoding and decoding schemes (Fig. 4B, detailed in Fig. S7). In simulations, a baseline GEO signal was first established using a fixed amount of carrier activator, and then a ramp input of data (fast activator) was introduced into the system to modulate the waveform. The resulting change in GEO frequency was estimated at every timepoint using Continuous Wavelet Transform (CWT), a time-frequency signal processing method^30,31^. Our simulations revealed that faster encoders led to larger changes in the FM response of the circuit, increasing encoding sensitivity; and different levels and rates of activation could be distinguished and decoded in the FM response of the circuit, demonstrating quantitative decoding (Fig. 4B).

Guided by these modeling results, we selected a GEO-FM circuit architecture that uses a slow *E. coli* activator/ATPase pair to generate a persistent barcoded carrier signal and the fast *P. damselae* activator for sensitive FM data encoding. We then used this design to build a circuit that streams transcriptional activity by placing the fast data encoder under a doxycycline-inducible TetON promoter (Fig. 5B)^32^. In parallel, we built a streaming circuit for fast timescale proteasome activity by connecting the GEO-FM encoder to the drug-controllable DHFR degron (Fig. 5C)^33^. Here, the fast activator is constitutively targeted for proteasomal destruction but can be rapidly stabilized upon addition of the small molecule TMP. To test the circuits, cells were imaged in the presence or absence of inducer at 0.25 fps for 18 hours using low intensity widefield fluorescence microscopy and instantaneous single-cell frequency over time was tracked using CWT (Fig. 4C, detailed in Fig. S10). The resulting single-cell data streams can be visualized directly as a continuous image timeseries, with pixel intensity set by signal amplitude and color-coded by frequency (Fig. 4D); or aggregated into stacks of single-cell tracks for comparison of dynamic responses either within or between conditions (Fig. 4F-G).

**Figure 5.**
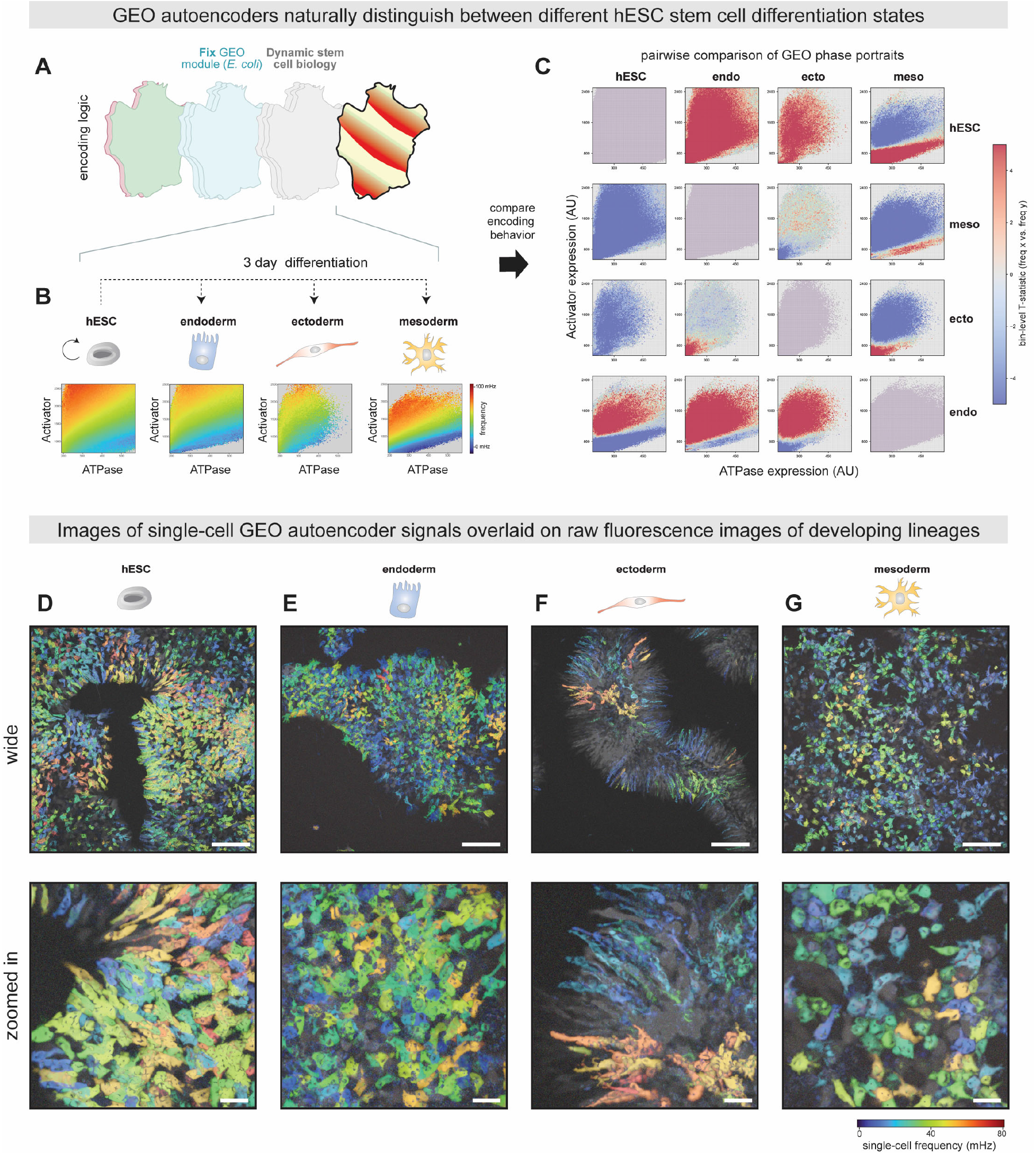
GEO autoencoders can distinguish human stem-cell differentiated states. (A) Conceptual schematic of a GEO autoencoder. When GEO biochemical components are fixed, the relationship between GEO expression levels and the resulting single-cell waveform can be influenced by changes in underlying cell physiology. This allows GEOs to act as an autoencoder of cell state and physiology. (B) GEO waveform encoding phase portraits for undifferentiated human embryonic stem cells (hESCs) and endoderm, ectoderm, and mesoderm germ layers three days post differentiation are shown as binned 2D histograms colored by the mean frequency at that location (see colorbar). (C) Pairwise comparisons of the GEO waveform encoding portraits from (B). Overlapping bins are colored by the Welch’s T-statistic obtained by comparing the row condition to the column condition. Red values indicate significantly faster oscillations at that location in expression space, while blue values indicate significantly slower oscillations (see colorbar). (D) Representative widefield (top) and zoomed in (bottom) images of the hESC control population from (B)-(C) at day 3, with single-cell GEO frequency overlaid on this image colored by frequency (see colorbar). Individual stem-cells generally appear as a single continuous color in this representation, corresponding to their associated unique GEO FM signal and barcode. Scale bars: top 200 um, bottom 40 um. (E-G) As in (D), but for the endoderm (E), ectoderm (F) and mesoderm (G) populations at three days post differentiation. The distinct single-cell morphologies associated with the different germ layer types are visible from the GEO FM barcoded signal. Scale bars: top 200 um, bottom 40 um.

For both circuits, single-cell GEO frequencies were stable in the absence of their small molecule inducer, showing little to no change over an 18-hour period. In contrast, addition of inducer generated potent single-cell GEO-FM responses that recapitulated the expected timescales of the regulatory dynamics at play^34^. For the transcriptional circuit, GEO-FM signal modulation took several hours to become detectable after induction and then increased robustly over the rest of the experiment (Fig. 4F). In contrast, GEO-FM signal modulation in the degradation circuit was detected within less than an hour and rapidly reached a new steady state, owing to the fast kinetics of encoder stabilization (Fig. 4G). Thus, GEO-FM circuits can successfully map dynamic changes in a specific cellular activity into GEO frequency modulation.

Because cells with different carrier signal barcodes will produce different FM responses as data encoder levels change, calibration is needed to interpret frequency changes and decode multiplexed data streams. To achieve this, we trained a machine-learning decoder model compatible with any circuit based on our GEO-FM architecture by measuring frequency across a broad range of randomized *E*.*coli* ATPase/activator carrier and *P. damselae* encoder expression levels. In addition to its general decoding utility, this model can also aid in the development of GEO-FM circuit design for specific applications by providing a landscape of the FM response space, revealing specific configurations with high dynamic range or high sensitivity.

Using this model, we mapped the single-cell GEO-FM responses of our transcriptional and degradation circuits back into their underlying changes in protein levels, allowing direct comparison of data streams from cells with different carrier signal barcodes (Fig. 4F-G). GEO-FM circuits can thus convert any existing intensity-based reporter scheme into an enhanced FM-barcoded single-cell data stream, measured in non-arbitrary units (ΔmHz) and robust against photobleaching and intensity fluctuations.

### GEO autoencoders can distinguish and classify human stem-cell differentiated states

The GEO streaming circuits developed thus far have leveraged the fact that a cell’s underlying carrier oscillation is stable within a consistent physiological state, enabling the deployment of engineered FM modulation circuits. However, cell physiology itself can serve as an additional waveform encoding layer. In this framework, GEOs function as *autoencoders* that translate large shifts in cellular state directly into changes in how GEO expression levels are encoded into waveform behavior (Fig. 5A).

To explore the utility and application of GEO autoencoders, we installed the *E. coli* GEO module in human embryonic stem cells (hESCs), a pluripotent cell type whose state can be maintained or directed toward diverse developmental fates. GEOs produced exceptionally clear signals across a broad range of expression levels, even within densely packed hESC colonies, generating sharp single-cell frequency beacons that revealed cell shape and spatial position with remarkable clarity (Movie S5 and Fig. 5D). To analyze waveform encoding behavior in this dense context, we modified our analysis pipeline to include only high-confidence pixels associated with a single unambiguous waveform. The resulting phase portraits followed the same general frequency-scaling relationships observed in immortalized cell lines (Fig. 5B), demonstrating that GEOs generate robust and quantitatively analyzable protein waves in a human stem cell context.

We next asked whether GEO waveform encoding behavior changed as hESCs differentiated into distinct germ-layer lineages. To this end, primed hESCs were induced to form endoderm, ectoderm, or mesoderm over a 72-hour period. The resulting populations displayed characteristic features of their respective developmental identities: mesoderm cells adopted irregular mesenchymal morphologies and organized into loose mesenchyme-like structures; ectoderm cells formed elongated tubular morphologies at colony boundaries; and endoderm cells exhibited the expected cobblestone-like appearance (Fig. 5D–G)^51,52^.

Importantly, GEO oscillations persisted in most differentiated cells, enabling direct comparison of waveform encoding behavior across developmental states. We therefore generated phase portraits from tens of thousands of measurements obtained from hESC, endoderm, ectoderm, and mesoderm populations 72 hours after differentiation (Fig. 5B). These data were aggregated into 2D histograms to facilitate statistical comparison of the frequencies associated with matched ATPase and activator expression levels across cell populations. Pairwise comparisons revealed striking differences in the structure of GEO waveform encoding behavior between developmental states (Fig. 5C). For example, hESCs generated faster oscillations than either endoderm or ectoderm across nearly all expression levels, whereas mesoderm cells occupied a broader frequency range despite exhibiting a narrower distribution of expression levels. Among differentiated populations, endoderm and ectoderm displayed the most similar encoding behavior, differing primarily at very low expression levels.

Together, these results demonstrate that GEO waveform encoding is strongly influenced by developmental state and cellular physiology. Differentiation into distinct germ layers systematically reshapes the relationship between GEO component expression and oscillatory behavior, creating characteristic waveform signatures that distinguish cell populations. As a result, GEO autoencoders provide a means of reading out cell identity directly from fluorescence dynamics, simultaneously reporting both developmental state and single-cell morphology within intact living stem-cell colonies.

### GEO visualization of stem-cell development and heterogeneity during differentiation

The ability of GEO autoencoder phase portraits to distinguish differentiated populations suggested that GEO waveform encoding could potentially be used to follow the trajectory of single-cell development within living populations. To explore this possibility, we generated GEO waveform-encoding phase portraits every 12 hours as primed stem cell populations either recovered into an hESC colony state or differentiated toward endoderm, ectoderm, or mesoderm fates (Fig. 6A, Fig. S11, Movie S6). Pairwise comparisons of GEO phase portraits across all conditions and timepoints revealed that while the four populations were initially similar, their waveform-encoding behavior progressively diverged during differentiation. Within a given lineage, GEO phase portraits separated by short intervals of time tended to be more similar than those separated by longer intervals, while populations undergoing different differentiation programs became increasingly distinct from one another (Fig. S11A).

**Figure 6.**
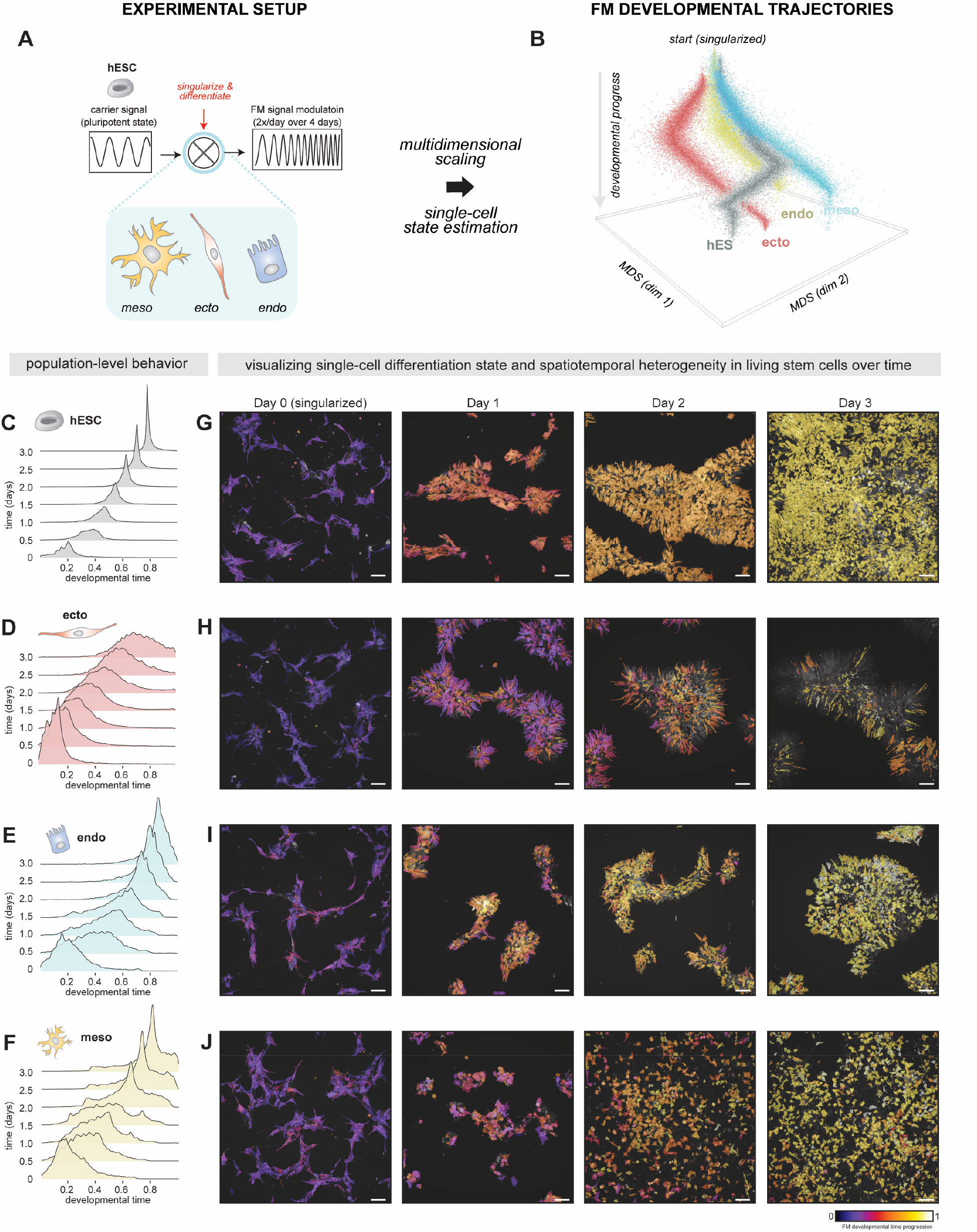
GEO autoencoders visualize single-cell development and heterogeneity during stem-cell differentiation. (A) Experimental calibration of GEO waveform encoding behavior across differentiating hESC populations over time allows for GEO autoencoder estimation of single-cell developmental progress. Waveform encoding portraits were collected for hESC, mesoderm, ectoderm and endoderm populations every 12 hours for 3 days across three biological replicates. Comparing all 2D histograms to one another (as in Fig. 5C) produces a matrix of all pairwise distances for dimensionality reduction and population-level trajectory estimation using multi-dimensional scaling (MDS). Single-cell ATPase, Activator, and frequency measurements are then used to estimate progress along these population-level developmental trajectories. See also Figure S11 and supplemental methods. (B) GEO-derived Waddington-like point-cloud visualization of single-cell developmental progress across hESC, endoderm, ectoderm, and mesoderm differentiation trajectories. Single-cell developmental time estimates were projected onto their respective positions on the population-level trajectory, displaced in proportion to uncertainty of the fit, and randomly jittered to improve visualization of observations. The X and Y axis correspond to MDS coordinates, and the Z axis corresponds to developmental progress. Beginning from a common starting point, the four populations diverge from one another over the course of development. (C-F) Kernel density estimate (KDE) plots showing the distribution of single-cell developmental time estimates present within the hESC (C), ectoderm (D), endoderm (E) and mesoderm (F) populations at each of the seven experimental timepoints. While hESCs display a smooth, largely homogeneous progression towards their recovered states, differentiated populations showed different patterns of single-cell heterogeneity over time. See also Figure S12. (G-J) Representative images of cell populations overlaid with color-coded GEO autoencoder estimates of single-cell developmental time (see colorbar). Consistent with (C-F), hESCs appear to progress homogeneously in both space and in time towards their colony state; while ectoderm, endoderm, and mesoderm show distinct spatiotemporal patterns of heterogeneity within the developing population structures. Scale bar: 100 um.

To organize and visualize these changes in waveform encoding, we aggregated bin-level differences between phase portraits to define a quantitative distance metric between any pair of populations. Pairwise distances across all differentiation conditions and timepoints generated a proximity matrix that was projected into a compact three-dimensional space using multidimensional scaling (MDS), where the first two dimensions were specified by dimensionality reduction and the third dimension represented developmental time. In this space, each differentiation condition traced a coherent trajectory from a common starting point, reminiscent of the metaphorical developmental landscapes originally envisioned by Waddington^53^. Importantly, the overall structure of this landscape was robust to different choices of bin-level statistics and distance metrics (Fig. S11B).

Having established population-level developmental trajectories, we next asked whether individual cells could be positioned along them. By comparing a cell’s ATPase and activator expression levels and GEO waveform properties to the expected behavior at each population-level timepoint, individual cells could be assigned to their most likely position along a developmental trajectory spanning the initial state (developmental time = 0) and terminal differentiated state (developmental time = 1) (Fig. 6B, Fig. S11; see Supplemental Methods).

This approach provides developmental-state estimates for every individual cell throughout the differentiation process, enabling direct visualization and quantification of developmental heterogeneity over time. To examine temporal changes in developmental heterogeneity, we generated kernel density estimates from these single-cell developmental-state assignments at each experimental timepoint (Fig. 6C–F). Recovery of hESCs back into colony growth occurred smoothly and homogeneously, with population standard deviation decreasing by more than 45% over the course of the experiment (Fig. 6C, Fig. S12). In contrast, each germ-layer differentiation trajectory exhibited a distinct pattern of developmental dispersion (Fig. 6D-F, Fig. S12). Ectoderm populations maintained broad developmental heterogeneity throughout the experiment, with population standard deviation increasing by more than 50% and many individual cells appearing to lag behind the average population trajectory (Fig. 6D). Endoderm and mesoderm populations both displayed a rapid increase in heterogeneity during the first 36 hours of differentiation but subsequently diverged: endoderm populations progressively converged toward a more homogeneous state, whereas mesoderm populations retained substantial heterogeneity through the end of the experiment (Fig. 6E,F). Notably, the temporal evolution of multiple heterogeneity metrics (standard deviation, skewness, and kurtosis) was highly reproducible across biological replicates, suggesting that these lineage-specific patterns of heterogeneity reflect underlying developmental programs associated with each developmental lineage (Fig. S12).

We next leveraged the unique ability of GEO autoencoders to project these single-cell developmental-state estimates directly onto images of living cell populations, enabling visualization of the spatial organization of developmental heterogeneity in both space and time (Fig. 6G–I). Consistent with our population-level analyses, hESC colonies exhibited minimal spatial or temporal heterogeneity. In contrast, distinct patterns of spatial organization emerged within differentiating populations. For example, ectoderm populations became enriched for elongated cells located near colony boundaries, and neighboring peripheral cells frequently exhibited similar developmental-state estimates. Likewise, the early increase in heterogeneity observed in endoderm and mesoderm populations initially appeared spatially well mixed but progressively resolved into patchy domains containing clusters of cells with similar developmental-state assignments. These observations suggest that developmental progression is not randomly distributed throughout a population but instead exhibits emergent spatial structure that evolves alongside differentiation. Thus, our approach provides a unique, and previously difficult to access, view into the spatial and temporal organization of single-cell behavior, coordination, and differentiation in a living multicellular developmental context.

Taken together, GEO autoencoders provide FM-barcoded readouts that enable continuous estimation of single-cell physiological state during complex developmental processes such as stem cell differentiation. Although the specific physiological changes responsible for the observed shifts in GEO encoding behavior remain incompletely understood, they manifest as readily measurable changes in waveform dynamics that can be used to track developmental progression in real time. We hypothesize that the differences in developmental progression and heterogeneity revealed by GEOs may arise from variation in the timing of differentiation initiation, local microenvironmental influences within developing structures, or interactions between these factors. More broadly, GEO autoencoders establish a framework for continuously mapping single-cell state within living multicellular systems. By providing longitudinal, spatially resolved measurements of cellular state without perturbing the underlying tissue, GEOs complement high-dimensional but destructive approaches such as single-cell RNA sequencing, as well as endpoint imaging methods based on immunostaining or in situ sequencing, opening new opportunities to study how cellular heterogeneity emerges, propagates, and organizes during development.

## Discussion

The GEO platform fulfills the demand for a high-performance biochemical analogue to the carrier signals that power modern telecommunications, unlocking a new paradigm for communicating live-cell biological information using frequency-modulated (FM) wave encoding and data transmission strategies. GEO-FM circuits that broadcast transcriptional and proteasomal dynamics demonstrate robust, multiplexed streaming of cellular information with high sensitivity and temporal resolution, while remaining resilient to photobleaching and systematic intensity fluctuations that often complicate long-term microscopy experiments. Likewise, GEO-autoencoder circuits allow shifts in cell physiology associated with changes in stem-cell development to be read out and decoded directly from changes in GEO wave dynamics.

By harnessing the evolutionary diversity of MinDE-family ATPase/activator pairs^35,54^, we greatly expand the ability to program and manipulate GEO waveforms in human cells. We show that ATPase and activator components can be treated as modular building blocks and establish design principles for combining, layering, and dynamically regulating GEO components to generate a broad range of circuit behaviors. Extensive characterization and modeling of GEO circuitry recover intuitive system parameters linked to the underlying biochemical properties of these modules, enabling scalable and predictable waveform engineering.

Once these biochemical encoding layers are established, a cell’s intrinsic physiology becomes the final determinant of GEO waveform. This principle was evident throughout our GEO variant library, where biochemical trends were conserved between human and yeast cells despite differences in their absolute waveforms. Building on this observation, we developed GEO autoencoder circuits that translate physiological variation into measurable changes in GEO dynamics. When deployed in human embryonic stem cells, these circuits revealed that differentiation into distinct germ layers systematically reshapes GEO encoding behavior. By calibrating these differentiation-dependent shifts, we were able to assign and visualize individual cells to developmental states, enabling direct visualization of single-cell identity within living stem cell colonies.

While GEO autoencoders effectively distinguish developmental states and cellular conditions, their broader applicability will depend on the degree of waveform divergence between the populations being compared. Because the physiological determinants of GEO waveform remain incompletely defined, GEO autoencoders are currently most effective for side-by-side comparisons of closely related cell populations. A more detailed understanding of how membrane composition, metabolism, and other features of cell physiology quantitatively alter GEO waveform in specific ways would further expand interpretability. Moreover, our current analyses leverage only a subset of the information contained within GEO patterns. Additional spatial features such as defect density, nodal architecture, wavelength, and other emergent pattern characteristics may provide substantial discriminatory power. Integrating these complementary GEO-derived features could ultimately enable GEOs to function as a live-cell analogue of Cell Painting^55^, in which complex cellular states are represented not by a static fingerprint but by a dynamic, multidimensional GEO feature profile.

More broadly, our approach of engineering synthetic circuits to manipulate the spatiotemporal dynamics of fluorescent proteins represents an alternative strategy for expanding the information content of fluorescence imaging. Rather than engineering new photophysical properties directly into fluorescent proteins, we use biochemical circuitry to encode information into how fluorescence evolves over space and time. This strategy complements decades of fluorescent protein engineering that have transformed biological imaging by modifying chromophore environments to generate new colors, alter excitation and emission properties, and introduce photo-switching or photo-conversion behaviors.^36–38^. These advances have dramatically expanded the experimental toolkit available to biologists and remain indispensable for visualizing cellular structures and molecular processes. However, spectral overlaps among optimized fluorescent protein combinations places practical limits on the number of simultaneously resolvable imaging channels ^39,40^. FLIM methods or FPs with distinct photo-switching times can exploit the timescale of an exponentially decaying fluorescent signal to further increase the number of usable channels^41–44^, but presently have palettes limited by our ability to understand and engineer the electronic state transitions at play.

GEOs extend this broader effort through a fundamentally different mechanism. Instead of encoding information through chromophore chemistry, GEOs use synthetic biochemical circuitry^45–50^ to couple a cell’s unique identity and internal state to the stream of emitted photons over time. This creates a rich set of imaging channels that are distinguishable by waveform dynamics and temporal frequency rather than color alone, enabling multiplexing and data-encoding applications using whichever fluorescent proteins are best suited for a particular biological system or experimental context. While engineered FPs remain essential for imaging subcellular structures, tracking molecular interactions, and capturing ultrafast signaling events, GEOs provide a complementary modality in which fluorescence serves as a dynamically programmable medium for communicating cellular information across biologically relevant timescales. By transforming fluorescence from a passive reporter into an actively programmable communication medium, the GEO platform provides a powerful high-performance data-encoding system that facilitates the acquisition and analysis of single-cell dynamics in real time, unlocking previously inaccessible FM data-encoding paradigms for continuous single-cell analysis.

## Acknowledgements

We thank members of the Coyle Lab, A. Weeks, W. Bement, W. Lim, and K. Roybal for advice, helpful discussions, and critical reading of the manuscript.

## Funding

This work was supported by a David and Lucille Packard Fellowship for Science and Engineering (SMC) and NIH New Innovator award 1DP2GM154329-01 (SMC). RR was supported in part by the National Institute of General Medical Sciences of the National Institutes of Health under Award Number T32GM008505 (Chemistry–Biology Interface Training Program). DB was supported in part by a Biotechnology Training Program NIH 5 T32 TGM135066A.

## Author contributions

Conceptualization RR, TMG and SMC.; methodology RR, TMG, JP, XZ, DB, EZWW, SMC; Investigation RR, TMG, JP, XZ, DB, EZWW, SMC.; writing – original draft, RR, TMG, JP, SMC; writing – review & editing RR, TMG, JP, XZ, DB, EZWW, SMC; funding acquisition, SMC; resources SMC.

## Competing interests

A provisional patent application has been filed by the University of Wisconsin and the Wisconsin Alumni Research Foundation related to this work.

## Notes

### Competing Interest Statement

The authors have declared no competing interest.

### Summary of Updates

Significant revision to the overall text and broader framing of the GEO technology. Added new stem-cell differentiation experiments and GEO autoencoder data (Figures 5 and 6). Added new supplemental materials (Movies S1, S5, S5; Figure S11-12).

