## Supplemental Material for "Single-cell FM streaming using genetically encoded protein oscillators"

**This PDF file contains:**

Materials and Methods.
Supplemental Figures S1-S12
Supplemental Movie Legends

**Materials and Methods.**

**Gene synthesis and cloning**

Genes encoding evolutionary variants of the MinD ATPase and MinE activator were obtained by gene synthesis as eBlocks Gene Fragments (Integrated DNA Technologies). Genes were codon optimized for expression in human cells using Benchling's website tool. MinD and MinE fluorescent protein fusions were generated by Gibson homology cloning into pLV-EF1a lentiviral transfer plasmids (addgene #85132); sequences were verified before use.

TetON MinE-BFP constructs were generated by subcloning into the pCW57.1 backbone (addgene #41393). The ccdB toxin gene was removed from the plasmid by standard molecular biology techniques prior to subcloning.

Gene encoding DHFR was obtained as gBlocks (Integrated DNA Technologies). DHFR C-terminal fusion to MinE-BFP was generated by Gibson homology cloning.

**Lentiviral transduction of mammalian cell lines**

Pantropic VSV-G pseudotyped lentivirus was produced by transfecting 293T cells (ATCC CRL-3216) with a pLV-EF1a transgene expression vector and the viral packaging plasmids pMD2.G (addgene #12259) and psPAX2 (addgene #12260) using Fugene 4K (Promega #E5911). Viral production was performed in 6-well tissue culture treated plates (Corning 3335). After 72 hours, viral supernatant was harvested, filtered using 0.45µM PES syringe filter, and added to mammalian cell lines with 2µg/ml Polybrene transfection reagent (Sigma TR-1003-G). When necessary, viral supernatant was concentrated by centrifugal filtration in a 100k MWCO PES Spin-X UF 20 Concentrator (Corning 431491) at 3214xg. Mammalian cells were exposed to the virus for 24 hr. After 5 days, cells were sorted on a FACSAria cell sorter (BD Biosciences) based on fluorescent protein expression levels. Cells were sorted to achieve diverse expression levels. Mammalian cells were expanded for 7 days before use in microscopy experiments. When necessary, cells were expanded without sorting and imaged immediately.

**Cancer cell lines culturing protocols**

Mammalian cells were cultured to a confluency of 60-80% in Dulbecco′s Modified Eagle′s Medium (DMEM) - high glucose (Sigma D6429) supplemented with 10% FBS (Cytiva Life Sciences SH30396.03) and 1% Penicillin-Streptomycin (ThermoFisher 15140122). At each passage, adherent cells were washed with PBS (at 37 °C) and TrypLE (ThermoFisher Scientific 12604021) was added to detach the cells from the flask surface. Flasks were incubated at 37 °C until the cells detached, typically 5 to 10 min. Fresh culture medium was added to quench the TrypLE and cells were resuspended and plated in new flasks and in fresh culture medium. The following cancer cell lines were purchased from the indicated vendors and cultured in the indicated media: 293T (ATCC CRL-3216) in DMEM and U-2 OS (HTB-96) in DMEM. All cell cultures were maintained in an incubator at 37 °C with 5% $CO_2$ and humidity.

**High-throughput screening of GEOs in mammalian cells**

293T cells were plated on 24-well glass-like polymer bottom plates (Cellvis P24-1.5P) to a confluency of 60-80% in FluoroBrite™ DMEM. The next day, cells were co-transfected with separate pLV-EF1a plasmids harboring the GEO ATPase and activator using Fugene 4K. Cells were imaged immediately the following day.

**Engineering GEO circuits in mammalian cells**
Cell lines expressing GEO circuits were generated for U-2 OS. Lentiviral particles for each genetic construct were generated separately, then pooled, filtered to remove cellular debris, and concentrated to a final volume of 2 ml before addition. Constructs were expressed under the control of an EF1a promoter. Cells were sorted on a FACSAria cell sorter (BD Biosciences) based on ATPase and activator expression levels and subsequently expanded and frozen at low passage. Engineered cell lines were sub-cultured until cultures reached passage 35 at which point fresh cells were thawed. Cell lines with multi-part GEO circuits were generated similarly or through lentiviral transduction of a single construct into an existing engineered cell line.

**Imaging GEO circuits in mammalian cells**
Cells were plated on glass bottom plates (Cellvis P06-1.5H-N) to a confluency of 60-80% in FluoroBrite™ DMEM (ThermoFisher A1896701). After 1 day, cells were imaged on a Nikon Ti-Eclipse in a Tokai Hit stage-top incubator: 37°C and 5% $CO_2$ conditions. Multi-channel red and green fluorescence images were collected at 1 FPS for 6 minutes. Multi-channel red, green, and blue fluorescence images were collected at 0.5 FPS for 6 minutes. For long continuous imaging experiments, images were collected at 0.25 FPS for 18 hours. For experiments that required drug addition, cells were incubated on the stage-top for 10 mins, before imaging.

**High-throughput screening of GEOs in yeast cells**
MinD ATPase (pTef1-mCherry-MinD-tAdh1) and MinE activator (pTef1-MinE-EGFP-tADH1) constructs were prepared as Cen/ars plasmids with leucine and histidine auxotrophic markers, respectively. Plasmids were transformed into *S. cerevisiae* strains BY4741 (*MAT***a** *his3*Δ1 *leu2*Δ0 *met15*Δ0 *ura3*Δ0) for MinD and BY4742 (*MAT*α his3Δ1 *leu2*Δ0 *lys2*Δ0 *ura3*Δ0) for MinE using the standard LiOAc/ssDNA transformation protocol. These transformed strains were plated in stripes on single selective plates, then replica plated onto YPD plates to perform the mating cross, then onto double selection plates (-his -leu) to select diploid cells carrying both plasmids.

Yeast cells were cultured in low fluorescence media (LFM). LFM was made consisting of 0.17% Yeast Nitrogen Base without Ammonium Sulfate, Folic Acid, or Riboflavin (MP114030512, Thermo Fisher Scientific), 0.5% Ammonium Sulfate, 0.2% appropriate amino acid supplement (with auxotrophic dropouts to enforce selection), where individual amino acids concentrations are defined in Yeast Synthetic Drop-out Media Supplements (Sigma-Aldrich), and 2% Glucose. Cultures were transferred to ConA-coated 96-well glass plate (Cellvis P96-1.5H-N) for imaging and were imaged on a Nikon Ti-Eclipse in a Tokai Hit stage-top incubator: 30°C conditions.

**Bioinformatic and evolutionary analysis of MinDE ATPase/activator superfamily**
AlphaFold2 using MMseqs2 was used to generate a protein structure prediction for the *E. coli* MinDE ATPase/activator dimer complex. The resulting MSA generated by paired MMseqs2 search, consisting of 4595 paired MinDE sequences, was used for downstream

bioinformatic and evolutionary analysis by RAxML (*antonellilab.github.io/raxmlGUI/).* Duplicate sequences in this search were removed, resulting in 4585 unique MinD ATPase and 4331 unique MinE activator sequences. These sequences were realigned using the ClustalW algorithm, then used to build phylogenetic trees using the JTT matrix for maximum likelihood with rapid bootstrap (100 BS) with gamma substitution rates for tree construction. The phylogenetic trees were visualized using iTOL (*itol.embl.de*). ConSurf analysis was performed using the protein structure prediction for the *E. coli* MinDE ATPase/activator dimer complex and the MinD ATPase and MinE activator MSAs and phylogenetic trees (*consurf.tau.ac.il*). ConSurf conservation scores were mapped onto the structure prediction. Pairwise distance between amino acid sequences were calculated as the patristic distances between pairs of sequences using the JTT-matrix-based model in MEGA (*megasoftware.net*).

**General image processing and GEO waveform analysis**

Waveform analysis of GEO behavior across cell lines (293T, U2OS, yeast) was performed using frequency-domain image analysis pipelines we established previously (Rajasekaran et. al. 2024), with the core tools available at `https://github.com/CoyleLab-UW-Madison/cellstream`. Briefly, basic image processing (cropping, background subtraction, and kymograph generation) was performed using Fiji ImageJ. The resulting image stacks were mapped from the time domain to the frequency domain using Fast Fourier Transform (FFT) on each pixel's intensity timecourse. This generates an "image FFT" in which each slice in the image stack corresponds to a different frequency, and the pixel value represents the complex-valued FFT result at that frequency slice. Taking the angle or magnitude of the complex valued FFT produces an image-level phase-field or power spectrum respectively.

For single cell analysis and reconstruction of waveform encoding phase portraits, cells were segmented in the frequency domain using an intensity threshold on each slice of the image power spectrum. The resulting masks are used to extract an average for that cell's fluorescence intensity in the MinE and MinD channels and the associated amplitude and frequency of its GEO waveform. This generates hundreds to thousands of single-cell measurements that can be used for downstream phase portrait construction or phenomenological modeling, as described below.

**Phenomenological models of GEO waveforms**

We constructed simple phenomenological models to gain insight into how specific GEO expression levels generate specific frequency and amplitude in mammalian cells. Note that the models presented in this paper are not intended to be a molecularly detailed description of reaction-diffusion dynamics rooted in thermodynamic and kinetic parameters. Instead, the goal of the model presented is to capture the minimal features of interest that we want to tune/control – e.g., how [Activator] to [ATPase] ratio scales with frequency, maximum frequency, how [ATPase] level scales with amplitude, and oscillatory functional space – thus providing a general framework to develop quantitative intuition about those properties to guide comparison between different GEO variants. Below we provide a detailed description of the features and assumptions of the presented model (additional detail in Fig. S5-6).

*Frequency and amplitude modeling*

The frequency model is a function of [MinD ATPase] and [MinE activator] expression levels and is a hyperbolic function composed of the key parameters: maximum frequency ($f_{max}$) and the half-$f_{max}$ [ATPase] to [Activator] ratio ($K$). The $f_{max}$ describes the maximum theoretical frequency a specific GEO pair can generate. The $K$ parameter describes how [ATPase] to [Activator] ratio scales with frequency. The amplitude model is a function of [MinD ATPase] expression level and is linear function composed of a single parameter: amplitude modulator ($a$). This suggests the amplitude is reasonably approximated by [MinD ATPase] expression levels and a GEO pair specific amplitude scaling factor.

*Oscillatory functional space modeling*

We developed a theoretical oscillatory functional space model to explain the observed differences in the [ATPase] and [Activator] expression levels that produce oscillations for different GEO combinations. This model considers that GEO waveforms are not only reaction-diffusion patterns, but mechanical waves with energy. Generally, mechanical wave energy is proportional to the frequency and amplitude of a wave, therefore we approximate the wave energy of GEOs as the product of our frequency and amplitude models. Note that this is not an explicit energy term but rather a useful relative approximation using known measurable parameters. Qualitative inspection of this model suggested that areas in expression space where changes in GEO expression levels resulted in higher wave energy directly correlated with the functional expression space we observed. These regions can be quantified using the product of the normalized (min-max scaled from 0 to 1) partial derivatives for wave energy. This term was normalized to represent an approximate probability for observing oscillations in a cell at a particular GEO expression level.

**PDE-based models and simulations of GEO circuits**

Simulations were performed using the mammalian cell setting (MCS) MinDE model we previously described (additional detail in Fig. S7). Multi-component GEOs were simulated by extending this model to include an additional MinE activator species with the same interactions and the same rate constants, with the only difference being each activator's respective ATPase activation rate. These simulations were analyzed using previously described FFT strategies. GEO-FM circuits were simulated by stepwise addition of this second MinE activator species into simulation space at discrete defined simulation intervals. These GEO-FM circuit simulations were analyzed using previously described CWT strategies.

**GEO-FM encoder circuits and analysis**

DOX-inducible transcriptional and DHFR degron stabilization pure FM encoder circuits were generated either by placing C-terminally BFP-tagged *P. damselae* activator MinE under the control of a Tet-responsive element (TRE) together with co-expression of rtTA (reverse tetracycline-controlled transactivator), or by direct C-terminal fusion of DHFR to the BFP-tagged encoder activator. Constructs were delivered into cells via lentiviral transduction. Low-intensity time-series fluorescence imaging was performed at 0.25 fps for 18 h in the presence or absence of inducer (2 μg/mL doxycycline or 10 μM TMP, respectively).

Pixel-level continuous wavelet transform (CWT) waveforms were computed using a generalized Morse wavelet ($\gamma = 3$, $\beta = 60$). Instantaneous frequency at each time point was assigned to the scale corresponding to the maximum wavelet coefficient. Single-cell frequency trajectories were extracted using TrackMate applied to Cellpose-generated segmentation masks derived from fluorescence image series temporally averaged over 180 s intervals. Each segmentation mask was subsequently applied to all time points within the corresponding interval to obtain instantaneous single-cell frequency measurements.

To map oscillation frequency to protein expression levels, a machine learning-based calibration model was developed to characterize the GEO-FM response space using experimentally measurable parameters. Training data were generated from cells expressing ATPase (mCherry), carrier activator (GFP), and encoder activator (BFP) components from independently integrated lentiviral constructs, resulting in heterogeneous expression levels across the population. Corresponding oscillation frequencies were measured for each circuit configuration. For this, a stacked ensemble regression framework was implemented using Random Forest Regression and Gradient Boosting Regression as base learners, whose predictions were integrated using an Elastic Net meta-model. The model was trained to predict encoder activator expression levels from ATPase expression, carrier activator expression, and oscillation frequency. 80% of the dataset was used for training, and the remaining 20% was reserved for cross-validation. The final model achieved an $R^2$ value of 0.782.

To use this calibration model to estimate encoder abundance in individual cells, the measured ATPase and carrier activator fluorescence intensities were supplied as fixed inputs to the trained model while oscillation frequency was systematically varied to generate a model-derived standard curve relating frequency to predicted encoder expression. The resulting linear relationship yielded a slope parameter that served as a conversion factor between frequency and encoder abundance. This conversion factor was subsequently applied to CWT-derived frequency changes to infer corresponding changes in encoder expression levels over time.

**Stem cell culture, manipulation, and differentiation.**
Human embryonic stem cells (WA09-H9, WiCell) were maintained under feeder-free conditions in serum-free mTeSR Plus medium on Matrigel-coated tissue culture plates using standard culture methods. Cells were cultured at 37 °C with 5% $CO_2$ and medium was changed daily. Matrigel-coated plates were prepared according to the manufacturer's instructions using DMEM/F12 media at least 1h prior to cell seeding. Colonies were maintained at approximately 60–80% confluency and passaged every 3–4 days using Versene cell dissociation agent.

To generate *E. coli* GEO-expressing stem cells, cells were transduced with concentrated and purified lentiviral supernatants. Lentiviral supernatant was collected 72h following transfection of HEK293T cells and clarified prior to purification. Lentiviral particles were purified and concentrated using the Vivapure LentiSELECT 40 membrane chromatography system (VS-LVPQ040, Sartorius) according to the manufacturer's protocol. Briefly, viral supernatant was mixed with loading buffer and passed slowly through the Sartobind Q membrane adsorber to allow selective binding of VSV-G pseudotyped lentiviral particles. The membrane was subsequently washed with washing buffer to remove residual media components, contaminating proteins, and nucleic acids. Bound viral particles were then eluted using high-salt elution buffer and further concentrated and buffer-exchanged into DPBS using Vivaspin 20 ultrafiltration columns (100 kDa MWCO). Purified lentivirus was either used immediately or stored at −80 °C for downstream applications.

For stem cell transduction, purified lentivirus encoding MinD and MinE constructs was added directly to pre-plated cells at a 3:1 ratio (MinD:MinE). Viral supernatant was removed following incubation, and transgene expression was assessed the following day by fluorescence imaging.

To perform differentiation experiments, hESC colonies were first dissociated into single cells using 0.25% Trypsin-EDTA and seeded into ibiTreat µ-Plates at a density of approximately $8 \times 10^4$ cells/mL in mTeSR Plus medium supplemented with 10% CloneR. Cells were allowed to recover overnight prior to differentiation induction. Each well was subsequently treated with lineage-specific differentiation media from the STEMdiff Trilineage Differentiation Kit (Catalog #05230, STEMCELL Technologies) according to the manufacturer's protocol. During differentiation, media was refreshed every 12h. Single-cell oscillatory dynamics were imaged using a 10× objective for 3 min time-lapse acquisitions with sequential dual-channel imaging performed at 1 s intervals between channels for each field of view.

**Population and single-cell waveform analysis of stem-cell differentiation trajectories.**

Our general approach to analyzing single-cell state behavior of undifferentiated and differentiated stem-cell populations was as follows: 1) generate population-level phase portraits that map expression levels to their frequency statistics for each differentiation condition and timepoint; 2) apply a metric that captures the distance/similarity between any two phase portraits; 3) use multidimensional scaling (MDS) on all pairwise distances to project population-level phase portrait trajectories into a lower dimensional space; 4) estimate single-cell progression along these trajectories by comparing the observed expression levels and frequency for each individual to the statistics of each phase portrait within the trajectory; 5) visualize these states by mapping the associated single-cell metrics onto their associated segmentation contours in the actual imaging data. Below we outline each step of this pipeline in greater detail.

*Population-level analyses.*

Because we found that stem cell populations were generally dense and often spatially overlapping, we increased the stringency of our frequency-domain analysis pipelines for more robust mapping of pixel-level GEO expression to waveform. Image timeseries were mapped to the frequency domain using FFT, and pixels were filtered to include only those for which both MinE and MinD showed an FFT peak with a signal-to-noise Z-score > 5 (quality filter); and only a single peak was detected in the FFT (overlap filter). Because in principle only a single pixel is required to estimate a cell's frequency, this restricts our analysis to the most robust signals in the data and reduces contaminating fluorescence from neighboring cells that can influence the expression level to waveform mapping. Using this approach, pixel-level data from each experimental condition were binned by MinE and MinD expression level and used to generate 2D histograms that capture phase portrait statistics (mean, standard deviation, and observation count) that describe waveform encoding behavior of the population on a given day in each condition.

To compare phase portraits to one another, we used the statistics of each 2D histogram bin to compute a bin-level Welch's T-statistic wherever portraits overlapped (see for example, Fig. 5 and Fig. S11). Intuitively, this visualizes how much faster or slower one condition runs at that location in expression space relative to another. Bin-level statistics were then aggregated into a single difference metric value by summing the squared T-statistics across the entire portrait. By computing a pairwise distance between all experimentally obtained phase portraits, we obtain a high-dimensional distance matrix and project it into a lower dimensional space for visualization of the differentiation trajectories using non-metric multi-dimensional scaling. This technique embeds the high-dimensional distances into a lower-dimensional (2D) coordinate system while preserving the relative dissimilarity between samples. A random seed value of 42 was used to ensure the reproducibility of the embedding.

To visualize the developmental state-space, we combined the spatial coordinates derived from MDS with the temporal component of the dataset. We projected these coordinates into a 3D visualization where the XY-plane represents the MDS dimensions and the Z-axis represents developmental time. Alternative statistics and distance metrics (Cohen's d, $\Delta$Freq) yielded similar overall trajectories but with different detailed arrangement within the MDS coordinate space (Fig. S11). These analyses were performed in python using `scikit-learn` and visualized in 3D space using the `seaborn` package and `pyQtGraph`.

*Single-cell remapping and visualization.*
While the data for each cell in these trajectory experiments was measured on a real-world day (lab reference frame), its biological state may deviate from the population average. The population-level waveform encoding portraits we collected and the relative distances between them provide anchor points by which we can estimating single-cell state progression along a given trajectory. This in turn allows us to resolve and visualize spatial and temporal heterogeneity in single-cell developmental progression within individual populations across different differentiation conditions.

To this end, we modeled a cell's progression along the population-level trajectory (developmental progression) as a weighted posterior estimate derived from its agreement with its corresponding population-level phase portrait at each timepoint. This approach is inspired by methods for trajectory inference in time-stamped single-cell RNA sequencing datasets, such as the Gaussian Process framework for Bayesian pseudotime[1,2]. For each high-quality oscillating pixel (as defined above) within a cell's segmentation mask, its associated developmental time estimate was given by summing over all days as:

$$t_{pseudo} = \frac{\sum(D \cdot L \cdot P)}{\sum(L \cdot P)}$$

Where D is the lab-reference frame day, L is the observation likelihood ($L = e^{-0.5 \cdot Z^2}$; $\mathrm{Z} = \frac{\mathrm{F_{obs}} - F_{\mathrm{bin}}}{\sigma_{\mathrm{bin}}}$), and P ($P = \exp\left(-0.5 \cdot \left(\frac{D - D_{collect}}{\tau}\right)^2\right)$) is a Gaussian prior centered on the observation's collection day to prevent unrealistic or non-biological jumps in time. The relaxation parameter τ was set to 1.5 days based on the overall population-level progression within the trajectories over time. From these pixel-level estimates, a cell's mean and standard deviation developmental progress estimate could be obtained from aggregation of all pixels within its segmentation mask.

Single-cell states were subsequently visualized with different types of analysis in three ways. First, we visualized these measurements along the population trajectory by projecting them onto the Waddington-like landscape produced from population-level MDS. For this, the population-level MDS trajectory was interpolated along a smoothed arc-length parameterization. A cell's developmental time metric was then be used to assign an idealized coordinate in the MDS space and randomly jittered to facilitate visualization of otherwise overlapping data points. To incorporate the uncertainty of our single-cell state estimation into the visualization, we applied a displacement vector to each observation whose magnitude was proportional to the standard deviation of that cell's psuedotime estimate. Together, this creates a dense cloud of single-cell observations around the underlying population level trajectory whose positions reflect both a cell's developmental progression estimate and its uncertainty.

To visualize the evolution of single-cell states over time across the different developmental timecourses, we generated sets of Kernel Density Estimation (KDE) ridge plots using the Joyplots package. To allow comparison of "start" and "end" across the different trajectories, raw psuedotime values were first min-max scaled to [0,1]. Using these distributions, we computed different statistical metrics of heterogeneity (standard deviation, CV, skewness, kurtosis) for each timepoint as well as the change in these metrics between timepoints (see Fig. S12).

Finally, single-cell state estimates were mapped back onto the resulting image contours to visualize the spatial organization of heterogeneity within the differentiating population (see, Fig. 6). Because the number of high-quality oscillating pixels was typically smaller than the number of pixels in the FFT-derived segmentation mask (which cleanly and effectively discriminates the boundaries of overlapping cells in the frequency domain), a cell's entire segmentation contour was flooded with its mean psuedotime value. In this way, the high quality pixel based measurements associated with a cell can be used to color the entire cell contour in the image by the estimate.

1. Tran, T.N., and Bader, G.D. (2013). Tempora: cell trajectory inference using time-series single-cell RNA sequencing data. *PLoS computational biology 16(9)* (2020): e1008205. https://doi.org/10.1371/journal.pcbi.1008205

2. Reid, J.E., Wernischm L. Pseudotime estimation: deconfounding single cell time series. *Bioinformatics* (32) 19, 2973-2980. https://doi.org/10.1093/bioinformatics/btw372

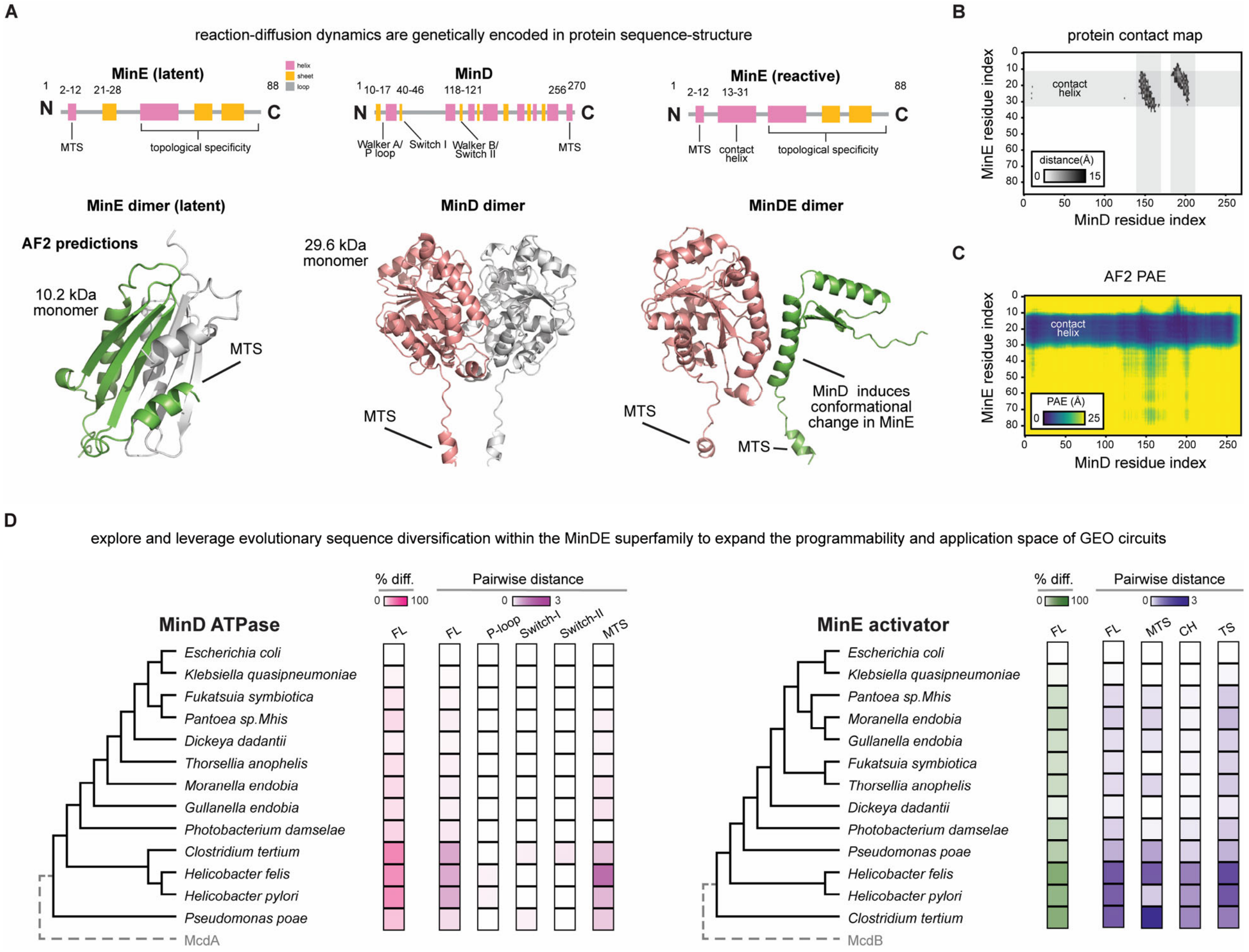


**Figure S1. Bioinformatic analysis of MinD ATPases and MinE activators reveals protein sequence diversity distributed across strucutre.**

(A) 3D molecular visualizations of AlphaFold2 structure predictions for the latent MinE homodimer, the MinD homodimer, and the MinDE heterodimer. Schematic views highlighting protein secondary structure predictions are shown: α-helical and β-strands structures are shown as pink and orange blocks; structural motifs and their respective amino acids are annotated. The corresponding secondary structure maps were attained from the AlphaFold2 structure predictions.

(B) The distance between two residues was calculated as Cα-Cα distance. Protein contact map highlighting close contacts between MinD and MinE in the AlphaFold2 MinDE dimer prediction is shown. A distance cutoff of 15Å was applied to the adjacency matrix.

(C) AlphaFold2 Predicted Aligned Error (PAE) for the MinDE dimer: low PAE between residues from different proteins indicates strong confidence in their relative positioning, suggesting an accurate spatial alignment. This supports the idea that MinDE interaction is primarily driven by the contact helix, with other regions of the proteins not significantly involved in the interaction.

(D) The percentage difference and pairwise distance were calculated for 13 species-matched MinDE protein sequences from various bacteria, using *E. coli* MinDE as the reference for these calculations. McdA ParA-type ATPase and its activator McdB were included as outgroups.

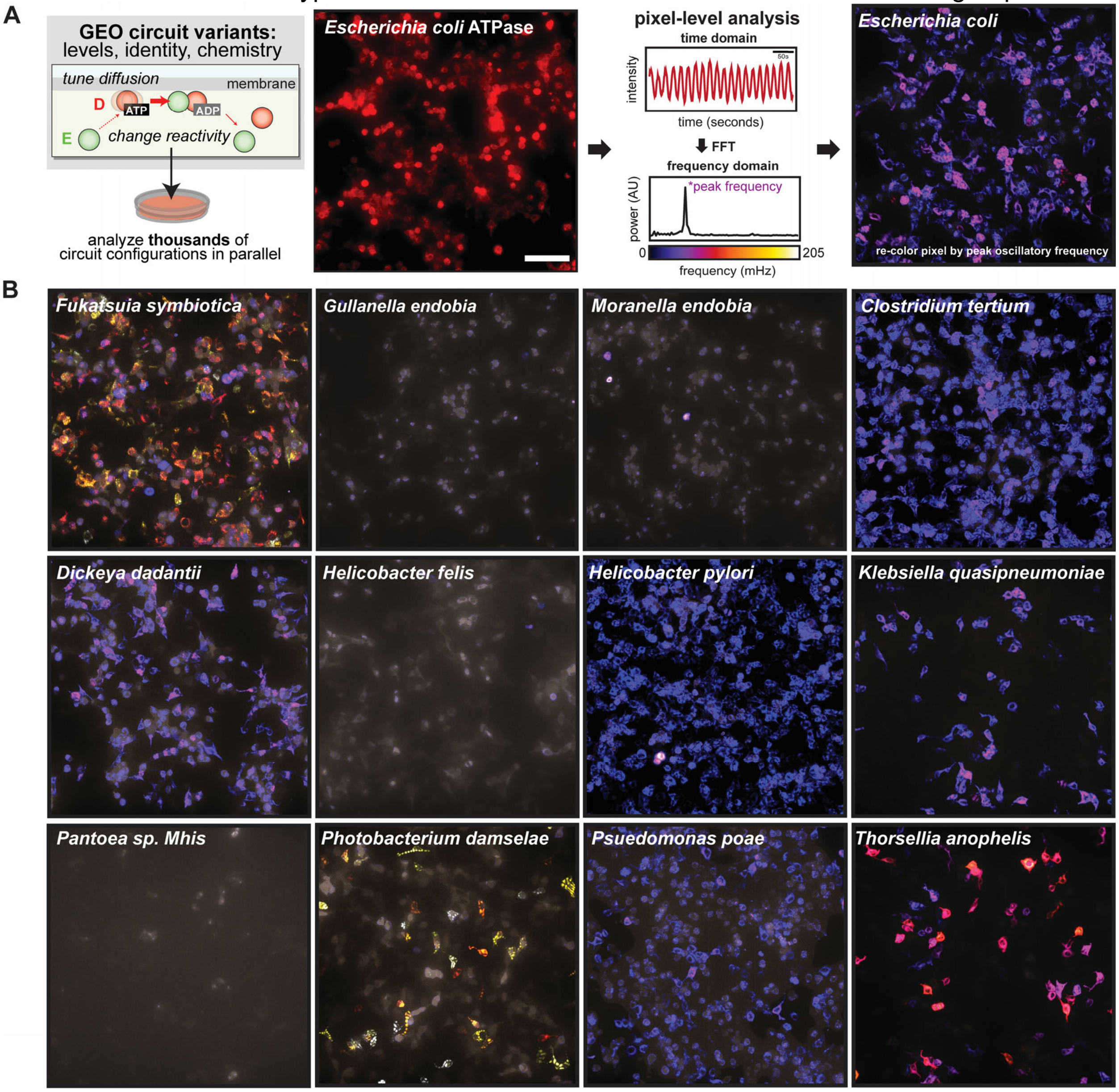


**Figure S2. Species-matched MinDE evolutionary variants generate a broad range of GEO circuit behaviors.**

(A) Schematic for assessing GEO functionality by imaging thousands of cell circuit configurations simultaneously. A representative low magnification widefield epifluorescence image of the *E. coli* GEO is displayed. FFT-based image processing transforms single-channel time series data into multiple frequency-defined channels, producing an image power spectrum that illustrates the distribution of oscillatory power across different frequencies. By false coloring each slice of the multi-channel image power spectrum according to peak oscillatory frequency, a single image is generated to visualize cells with different frequencies within the population. (scale bar: 100μm)

(B) Frequency re-colored images of species-matched GEO pairs.

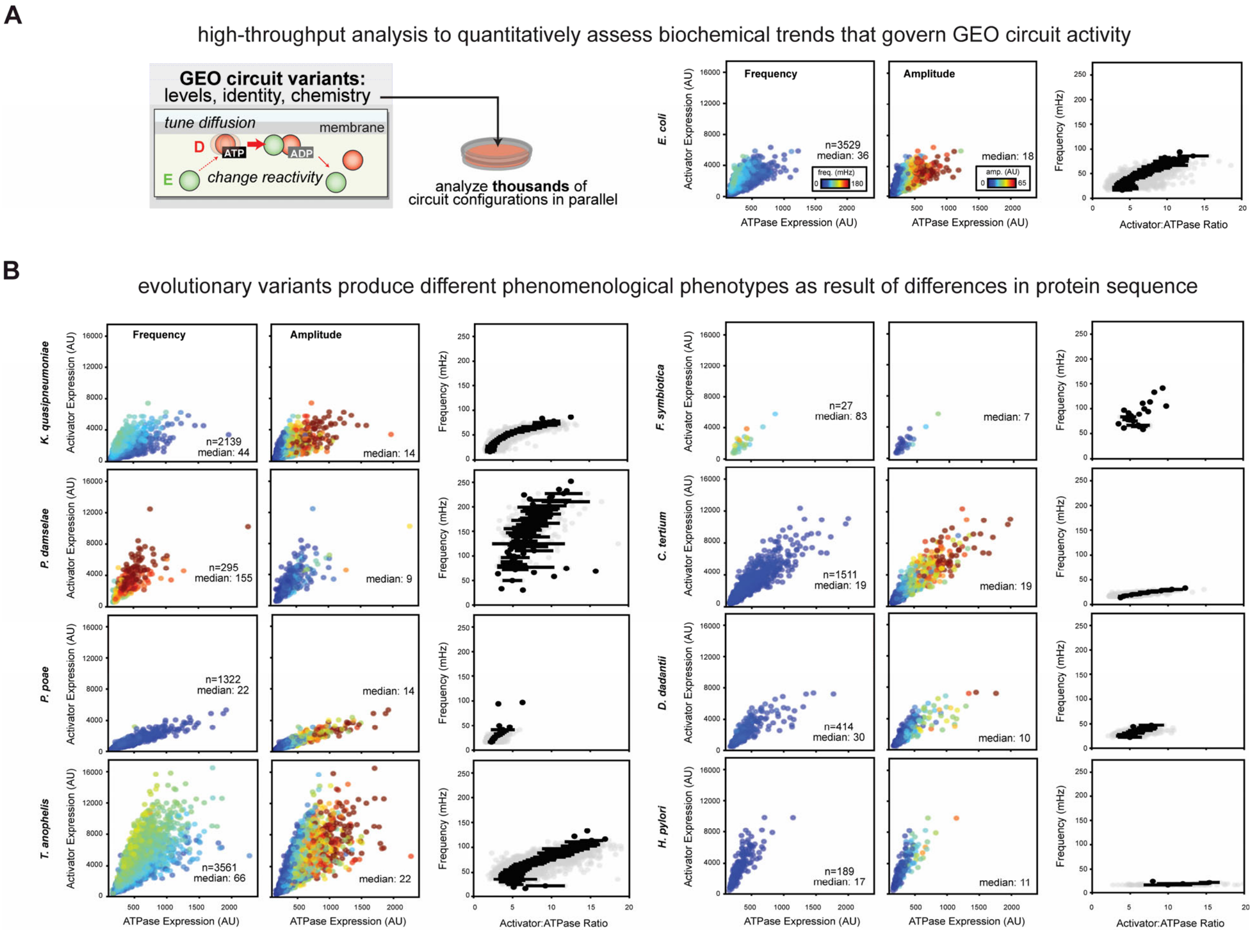


**Figure S3. Biochemical trends governing GEO circuit behaviors are diverse across species-matched MinDE evolutionary pairs.**

(A) High-throughput analysis of single-cell data from cells expressing different species-matched GEO pairs was performed to identify biochemical trends in circuit behavior. Low-magnification widefield epifluorescence images of mCherry-MinD (ATPase) and MinE-GFP (Activator) were captured. Frequency-domain image analysis was applied to spectrally separate and FM-barcode each cell. Standard OpenCV image processing was used to isolate FM-barcoded cells and extract frequency, amplitude, and mCherry/GFP fluorescence intensities (used as a proxy for protein expression level) for each cell. The single-cell data collected from many low-magnification widefield images were then aggregated to generate datasets for each species-matched GEO pair. Analysis of the *E. coli* GEO pair shows that frequency is governed by [MinE Activator] expression level, amplitude is governed by [MinD ATPase] expression level, and frequency is governed by the ratio of [Activator] to [ATPase]. Median value for frequency and amplitude across the population is indicated.

(B) The biochemical trends observed for the *E. coli* GEO are consistent across functional species-matched GEO pairs, suggesting a common set of biochemical principles that govern the behavior of GEO reaction-diffusion systems in mammalian cells. However, the scaling relationships of these trends varies among the functional species-matched GEO pairs tested, likely due to differences in the underlying protein biochemistry caused by variations in protein sequences.

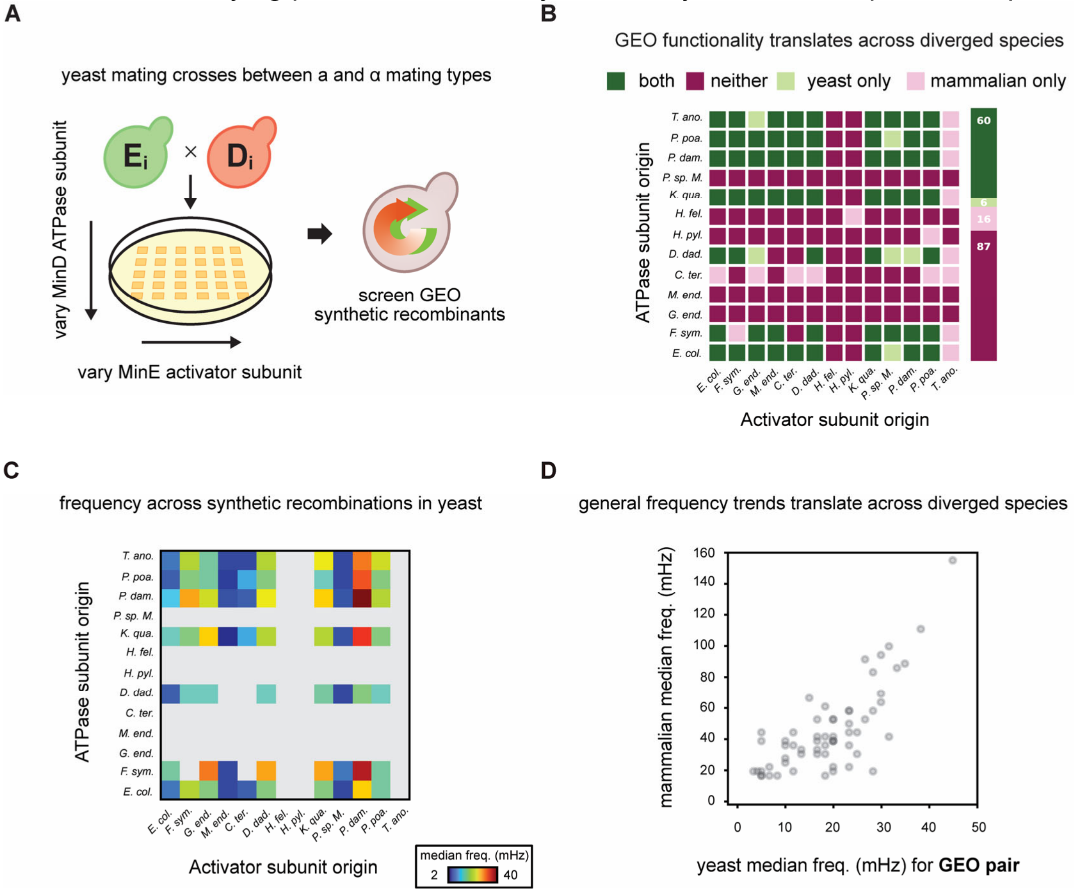


**Figure S4. Characterization of GEO circuits in budding yeast.**

(A) Schematic illustrating high-throughput GEO circuit screening in yeast. The process of combinatorial exploration of GEO circuits was accelerated through yeast mating crosses. Haploid yeast cells of mating types a and α expressing either the MinD ATPase or the MinE activator were prepared. A simple cross was performed to simultaneously generate all hybrid GEO combinations on a single plate.

(B) Diagram summarizing the shared functionality of GEO circuits in mammalian and yeast cells. Most GEO pairs are functional in both systems or neither. A small subset of GEO pairs show exclusive functionality in either yeast or mammalian cells. This indicates that the functionality of GEO pairs is context-dependent. While some GEO pairs may not function in the conditions tested here, it is possible that specific conditions could promote oscillations for these pairs.

(C) Heatmap showing the median population frequency of each GEO recombination pair in yeast, with non-functional GEOs represented in gray.

(D) Comparison of the median population frequency of each GEO recombination pair in both mammalian and yeast cells. The relative performance of GEO pairs is consistent across these two diverged species.

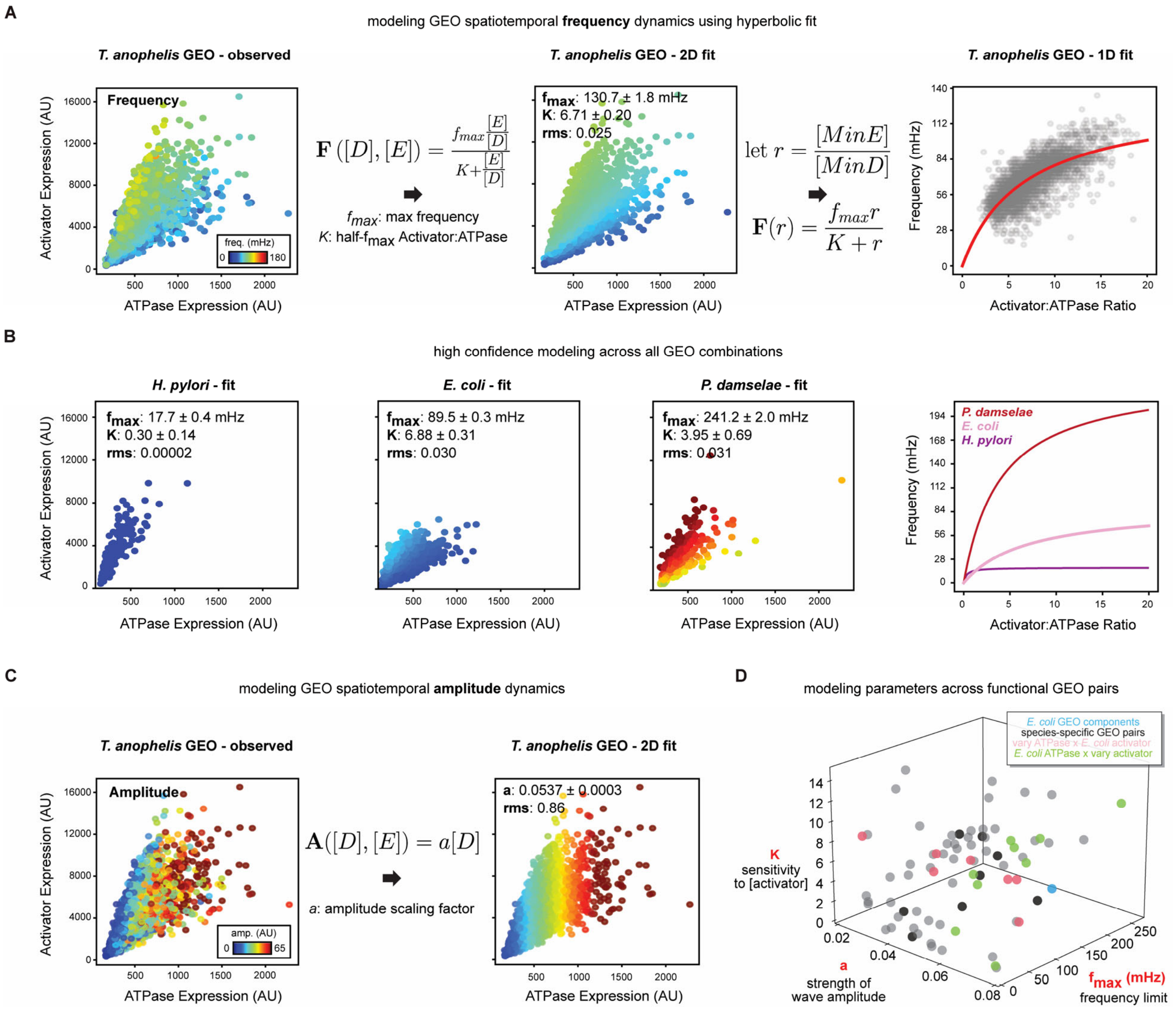


**Figure S5. Phenomenological modeling of GEO waveforms enables quantification and comparison of GEO behavior across hybrid pairs.**

Modeling of GEO dynamics to mathematically connect biochemistry with phenomenology was performed. Simple phenomenological models were constructed to better understand how specific GEO expression levels result in specific frequencies and amplitudes in mammalian cells.

(A) Representative frequency phase portrait for the *T. anophelis* GEO is shown. GEO frequency depends on [MinD ATPase] and [MinE activator] expression levels and hyperbolically scales with [MinE Activator] to [MinD ATPase] ratio. The model consists of two key parameters that intuitively describe important phenomenological features of oscillatory frequency: the $f_{max}$ represents the maximum theoretical frequency a specific GEO pair can generate, while the *K* parameter describes how the [ATPase] to [Activator] ratio influences frequency. A 2D fit with data points colored according to the model-predicted frequency is shown. A 1D fit, simplifying the [MinE Activator] to [MinD ATPase] ratio to a single term, *r*, is also shown, revealing how frequency scales with the [MinE Activator] to [MinD ATPase] ratio.

(B) Examples of the frequency model being applied to fit different species-matched GEO pairs.

(C) Representative amplitude phase portrait for the *T. anophelis* GEO is shown. GEO amplitude linearly scales with [MinD ATPase] expression. This suggests that amplitude is reasonably approximated by [MinD ATPase] expression level and a GEO pair specific amplitude scaling factor *a*. A 2D fit with data points colored by the model-predicted amplitude is shown.

(D) 3D scatterplot of $f_{max}$, *K,* and *a* for 76 functional GEO systems is presented.

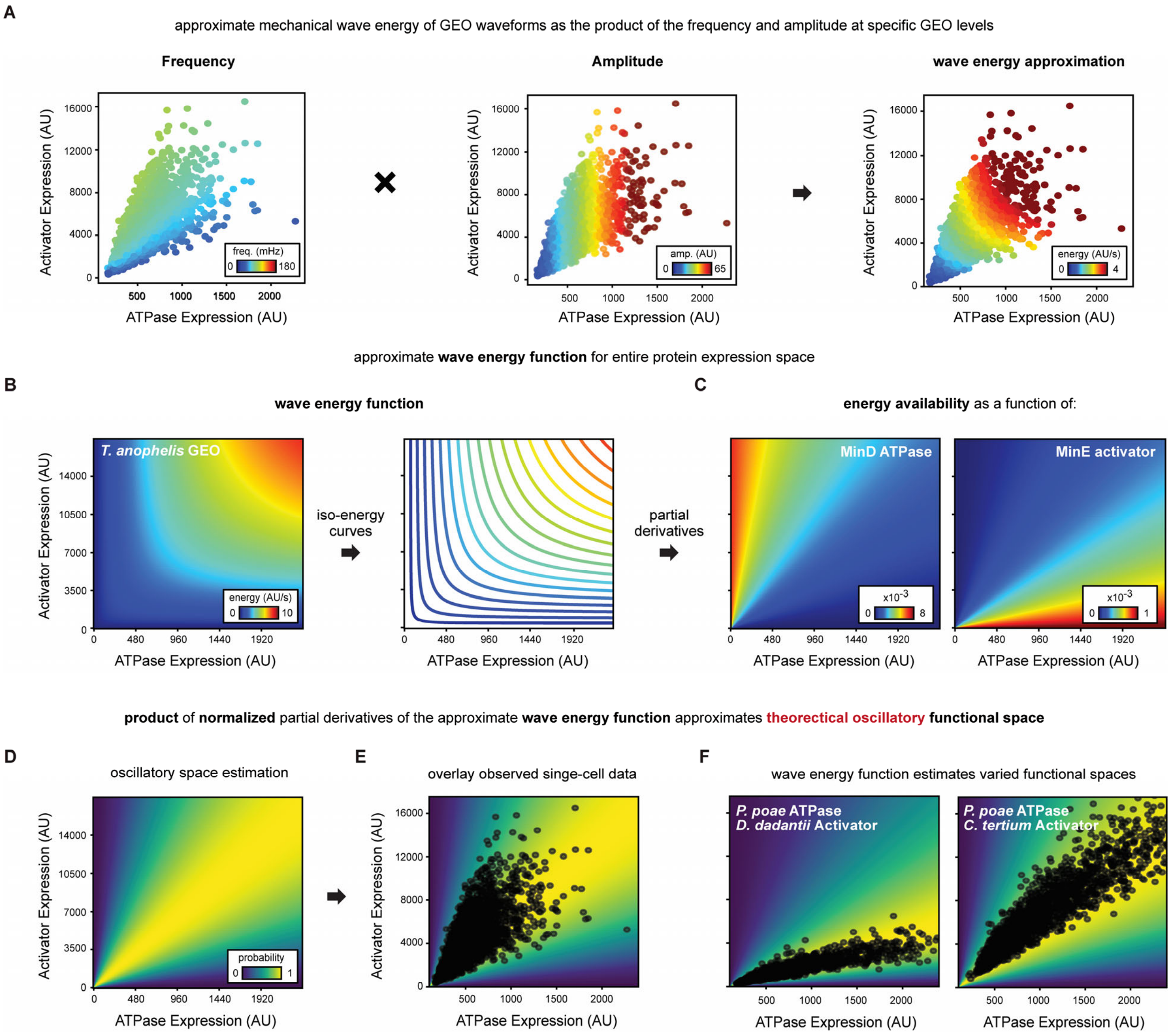


**Figure S6. Wave energy predicts regions of expression space that support oscillatory dynamics.**

Modeling of oscillatory functional space was performed to explain the observed variations in ATPase and activator expression levels that produce oscillations in different GEO combinations.

(A) Typically, mechanical wave energy is proportional to both the frequency and amplitude of a wave. Therefore, we approximate the wave energy of GEOs as the product of our frequency and amplitude models. Representative wave energy for the *T. anophelis* GEO is shown for regions of expression space where oscillating cells were experimentally observed.

(B) Wave energy for the *T. anophelis* GEO was calculated across the entire expression landscape. The resulting wave energy landscape reveals iso-energy lines, indicating that different expression levels can share the same wave energy. A qualitative inspection of this landscape, in conjunction with the experimental data, suggested that regions of expression space where changes in GEO expression levels result in higher wave energy correlate directly with the functional expression space observed. Thus, regions in expression space where changes in GEO expression levels do not result in higher wave energy represent areas that do not support oscillations.

(C) To quantify regions where changes in GEO expression levels do not result in higher wave energy, energy availability, defined as the partial derivatives of the wave energy function, was calculated. Regions of expression space with low energy availability indicate that changes to GEO levels in these areas do not significantly alter wave energy, marking these regions as unable to support oscillations.

(D) The energy availability for MinD and MinE were combined into one term by multiplying the normalized (min-max scaled from 0 to 1) MinD and MinE energy availability together. This combined energy availability term was then normalized (min-max scaled from 0 to 1) to approximate the theoretical oscillatory functional space. This model estimates the probability of observing functional oscillations in a cell with a given GEO expression level combination. The theoretical oscillatory functional space for the *T. anophelis* GEO is shown.

(E) The theoretical oscillatory functional space for *T. anophelis* GEO is shown with the expression levels of experimentally observed oscillating cells overlaid.

(F) Theoretical oscillatory functional space for the *P. poae* ATPase / *D. dadantii* activator and *P. poae* ATPase / *C. tertium* activator GEO pairs are shown, with the expression levels of experimentally observed oscillating cells for each respective pair overlaid.

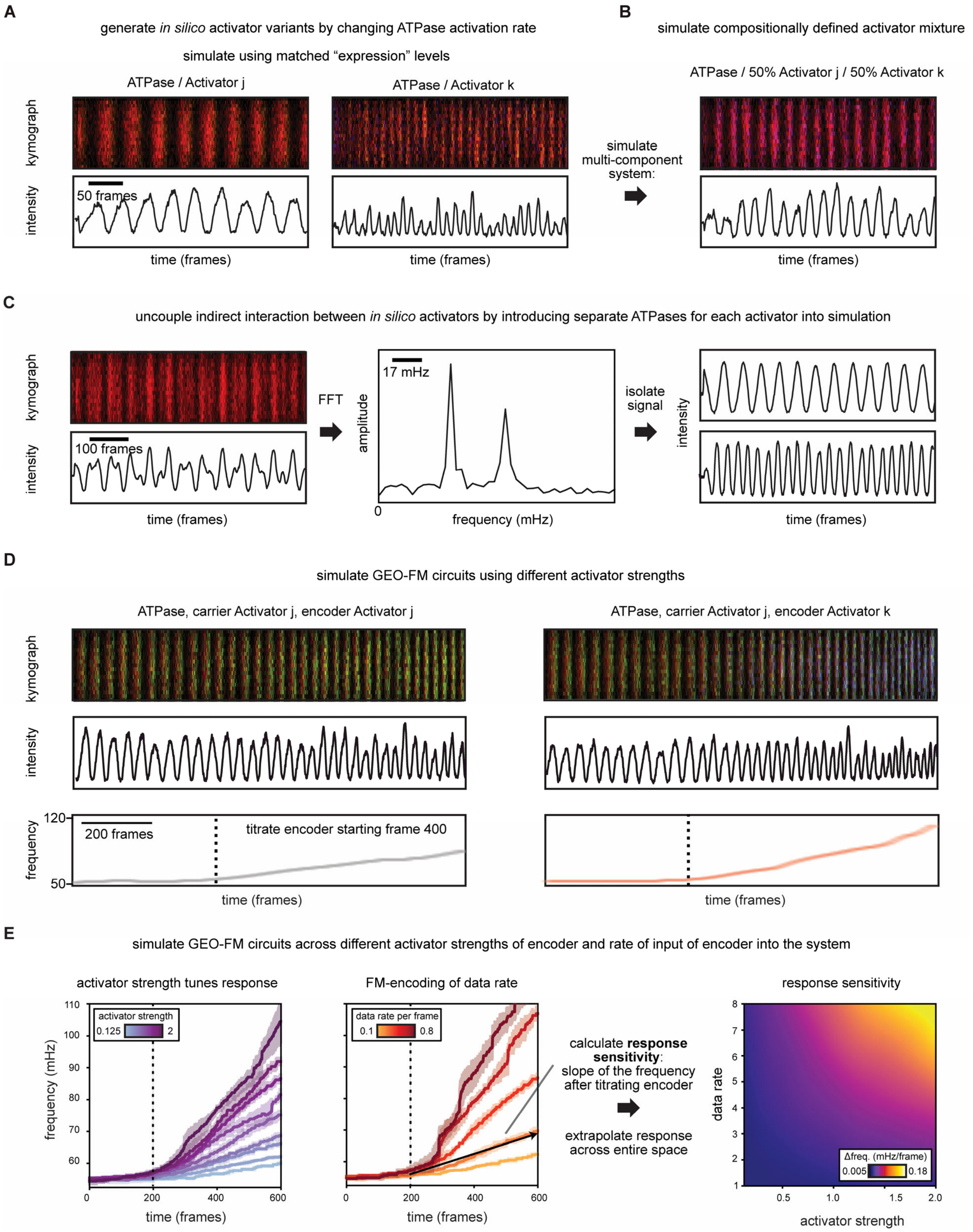


**Figure S7. Stochastic simulations of designed multi-component GEO circuits provide intuition into waveform behavior.**

(A) Representative kymographs and pixel-level timecourses from simulated microscopy images of single ATPase / single activator GEO circuits. Expression-level matched simulations using activators with two different ATPase activation rates are shown.

(B) Representative kymograph and pixel-level timecourse from simulated microscopy images of a multi-component GEO circuit with a compositionally defined activator mixture. Indirect coupling of MinE activator species through the shared MinD ATPase node results in a single waveform, in which the two MinE activator species oscillate in phase at a shared frequency.

(C) To test whether removing the indirect coupling would lead to separate waveforms, a distinct MinD ATPase species was introduced into the reaction scheme, and each respective MinDE pair was prevented from interacting explicitly. Representative kymograph and pixel-level timecourse from simulated microscopy images of this system are shown. Simulated microscopy images were generated by treating different MinD or MinE species as one. FFT-based signal processing was applied to the pixel-level data, revealing two distinct oscillatory frequencies in the simulation. FIR filters separated the independent signals generated by the separate GEO circuits.

(D) Examples of simulated GEO-FM circuit designs. A simulated carrier waveform composed of an ATPase and a carrier activator was given input data in the form of a ramped increase of an encoder activator. The simulated waveform was tracked to measure the GEO response, and CWT was used to extract the FM response. Simulations using encoder activators with two different ATPase activation rates are shown.

(E) Extended examples of GEO-FM circuit simulations with altered input data rates and encoder activator strengths are shown. Response sensitivity, calculated as the rate of frequency change over time in the response trajectory, was assessed for a set of data rates and encoder activator strengths. These sensitivities were then used to extrapolate the response sensitivity across the entire landscape.

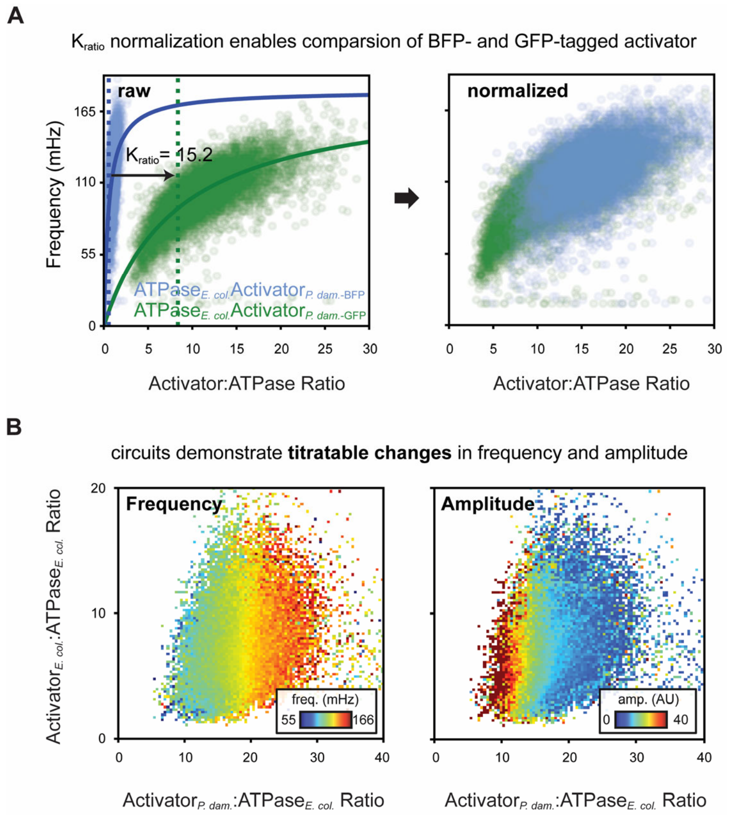


**Figure S8. Multi-dimensional phase portraits reveal biochemical trends that govern multi-component GEO circuit behavivor.**

(A) To directly compare and combine the levels of different activators tagged with distinct fluorescent proteins, a normalization strategy was developed to convert BFP expression level into approximate GFP expression level. This approach leverages frequency, a true measurement independent of fluorescence. Since the same GEO pair should yield the same frequency behavior regardless of the fluorescent proteins used for visualization, we tested the same GEO pair with the activator tagged either with GFP or BFP. These data were modeled as outlined in Fig. S5. We found that while both datasets shared the same $f_{max}$, their $K$ parameters differed significantly. This difference is expected, as frequency is independent of imaging parameters, and the [Activator] to [ATPase] scaling relationship is sensitive to imaging conditions and fluorescent proteins. Given that the same fluorescent protein was used for the ATPase, we hypothesized that the difference in the apparent $K$ value is solely due to the fluorescent protein attached to the activator. As a result, the ratio of the $K$ parameters can be interpreted as the ratio between the fluorescence outputs of a BFP-tagged and GFP-tagged protein. By using this ratio as a conversion factor, the levels of BFP-tagged and GFP-tagged activators in the same cell can be compared and combined.

(B) Multi-dimensional phase portrait of this multi-component GEO circuit shows that circuit behavior is governed by relative protein levels. An inverse relationship is observed between frequency and amplitude.

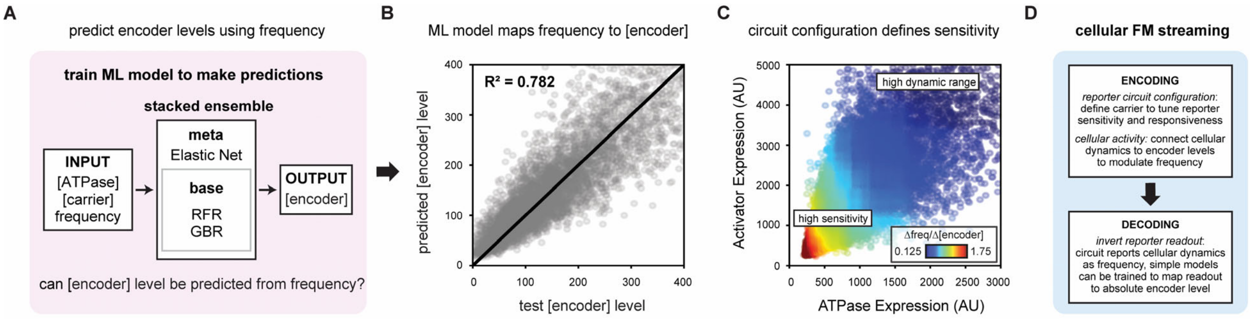


**Figure S9. Machine-learning model calibrates GEO-FM response to underlying biological data of interest.**

(A) Schematic of the machine-learning model used to calibrate GEO-FM response based on measurable parameters. A stacked ensemble machine-learning approach was employed, combining Random Forest Regression and Gradient Boost Regression as base models, whose predictions served as inputs for the Elastic Net meta-model. The model was trained using known ATPase (mCherry), carrier activator (GFP), and encoder activator expression levels (BFP), alongside the corresponding frequency for each circuit configuration. The model aims to predict encoder levels based on ATPase and carrier activator levels and the corresponding frequency. 80% of the data was used for training.

(B) Cross-validation of the model was conducted using 20% of the data. An R-squared value of 0.782 indicates that the model effectively captures the relationship between ATPase, carrier activator, encoder activator, and frequency, without overfitting.

(C) The machine-learning model was applied to assess the sensitivity of different ATPase and carrier activator configurations to changes in encoder levels. For each carrier signal configuration, the model predicted encoder levels across a small range of frequencies. The linear relationship between frequency and encoder level changes defines the circuit's sensitivity. Regions of ATPase and carrier expression space with high sensitivity require minor changes in encoder level to induce large frequency changes, while regions with low sensitivity require large changes to encoder level to achieve the same frequency shift.

(D) Overview of the GEO-FM paradigm for biological data transmission.

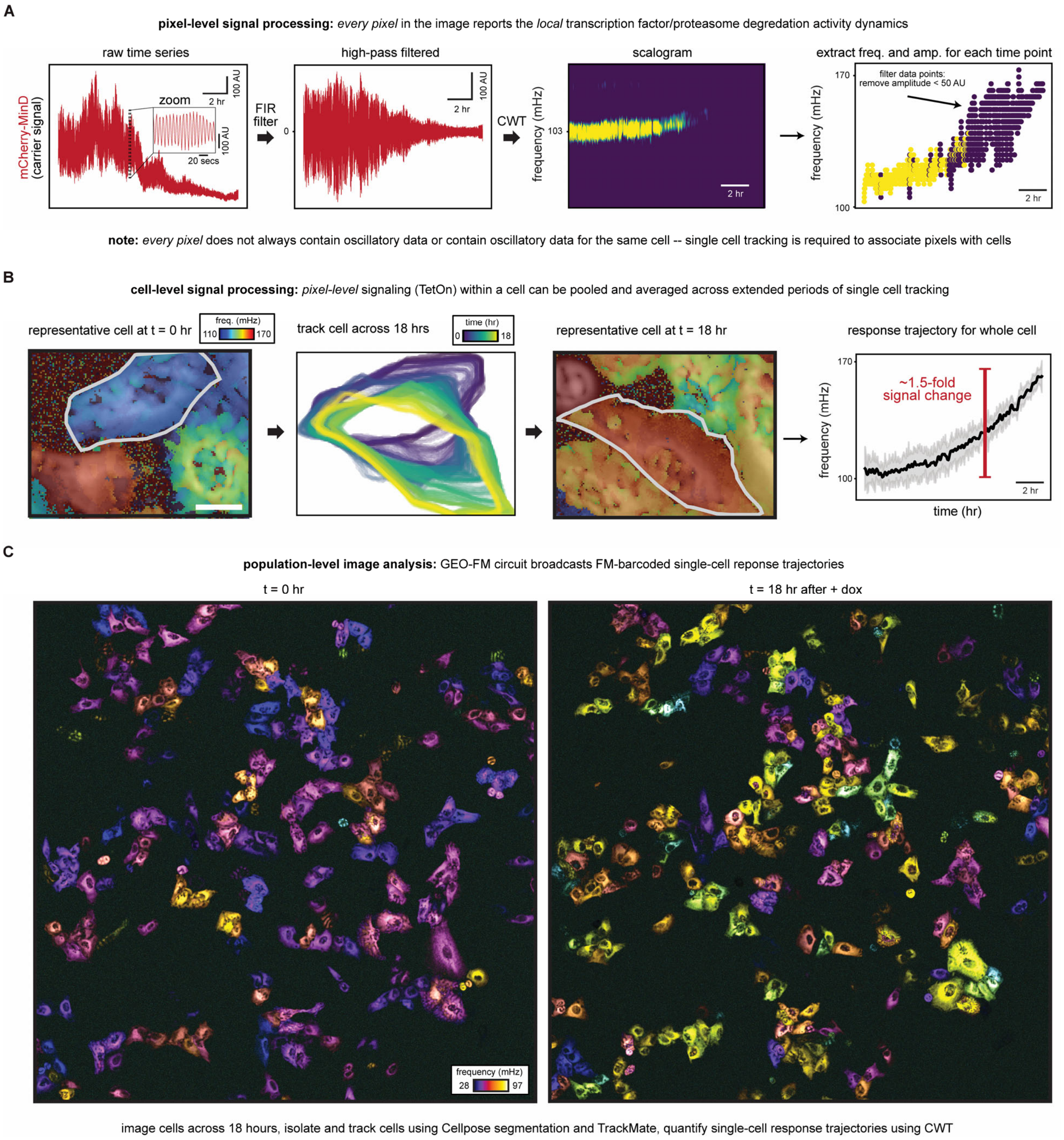


**Figure S10. Digital signal processing workflow for quantifying single-cell GEO-FM response.**

The GEO-FM circuits we present here enable the characterization and quantification of transcription factor activity or protein degradation dynamics at a single-pixel resolution, offering a detailed, localized readout of cell state behavior. Pixel-level data can be aggregated at the cellular level to generate single-cell response trajectories for transcription factor activity or protein degradation dynamics.

(A) At the single-pixel level, the raw fluorescence time series for mCherry-MinD (the carrier line) reflects the local GEO-FM circuit response. The DC components of this signal were eliminated using an FIR filter, resulting in zero-centered signals. The instantaneous frequency throughout the imaging experiment was then computed using the Continuous Wavelet Transform (CWT). This produces pixel-level scalograms reporting on frequency power content across times for different scales that each represent a unique frequency. The scale with the highest magnitude at a given time was assigned as the dominant oscillatory frequency for that pixel at that time, while the magnitude of that scale coefficient was assigned as the pixel's amplitude. The resulting frequency signal provides insight into the local signaling activity within the cell, showing a frequency increase over time. High-frequency noise is characterized by low amplitude and can be excluded during pixel-aggregation steps.

(B) To generate single-cell GEO-FM trajectories, the pixel-level data in (A) was grouped by all pixels belonging to a single cell, with cell boundaries identified for each frame using Cellpose segmentation masks. Pixels within the cell were filtered based on an amplitude threshold to exclude regions of high-frequency, low-amplitude background noise that might be included in the Cellpose masks. The frequency of these pixels was then averaged across each time point for the cell. A representative cell and example analysis is shown. Cell boundary tracks obtained using TrackMate are shown. Frequency-colored images at the start and end of the experiment for this cell highlight a significant shift in frequency. For this cell, the GEO-FM trajectory shows a 1.5-fold increase in frequency over the course of the imaging experiment.

(C) To extract single-cell trajectories from wide-field imaging of a population of cells, image-level CWT was applied to the MinD ATPase carrier signal. Representative frequency false-colored images of a population of cells undergoing a GEO-FM response are shown for both the start and end of the imaging. Cells can be isolated and tracked as outlined in (B), to quantify the individual single-cell response trajectories.

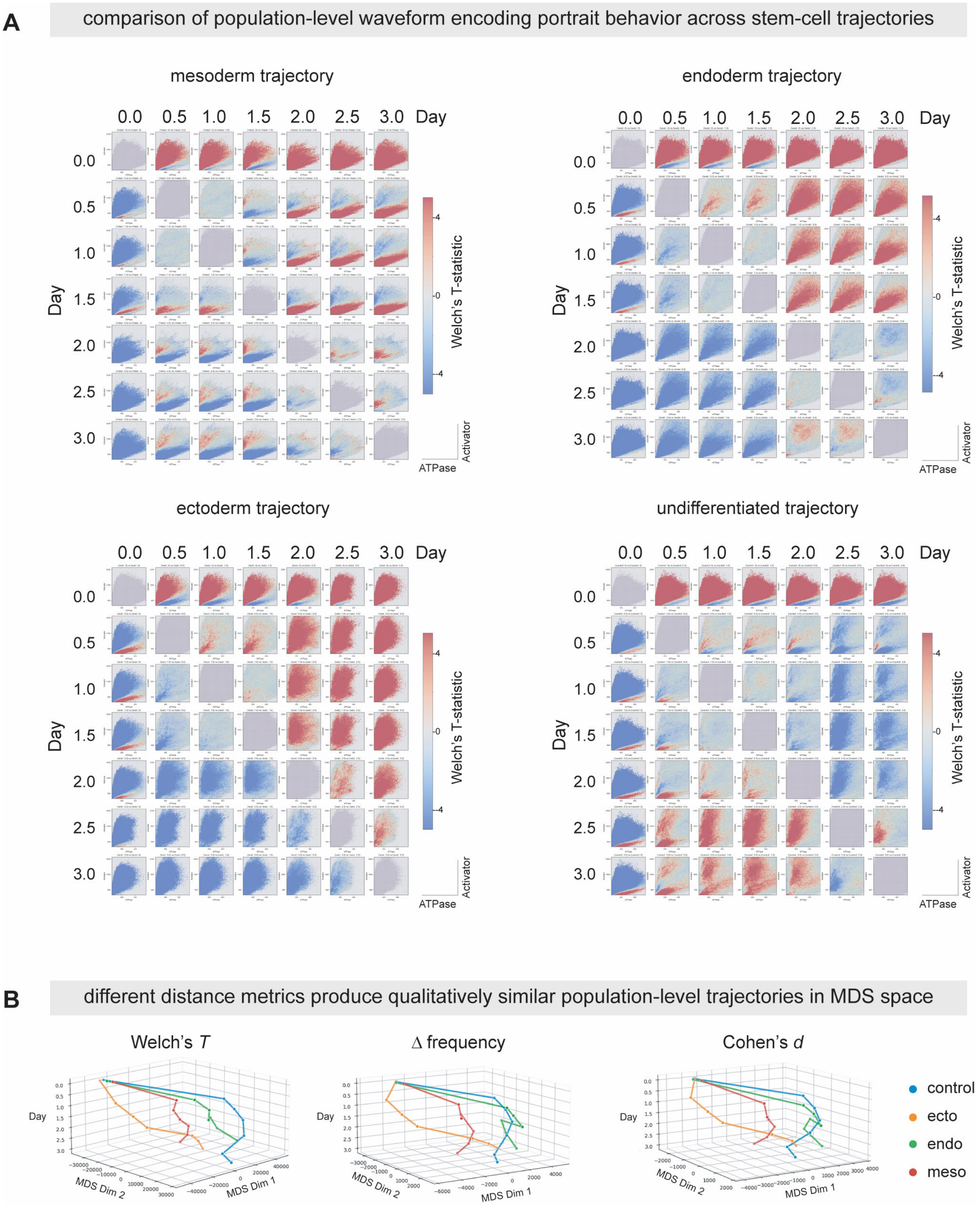
A
comparison of population-level waveform encoding portrait behavior across stem-cell trajectories
mesoderm trajectory
endoderm trajectory
ectoderm trajectory
undifferentiated trajectory
0.0
0.5
1.0
1.5
2.0
2.5
3.0
Day
Welch's T-statistic
4
0
-4
Activator
ATPase
B
different distance metrics produce qualitatively similar population-level trajectories in MDS space
Welch's T
Δ frequency
Cohen's d
Day
MDS Dim 1
MDS Dim 2
control
ecto
endo
meso

**Figure S11. Pairwise comparison of stem-cell waveform encoding portraits during differentiation and the effects of metrics on resulting MDS trajectories.**

(A) Comparison of waveform encoding behavior across control, endo, ecto, and meso differentiation trajectories over 3 days. 2D histograms corresponding to a cell's (E,D) to frequency encoding profile were compared at a bin-to-bin level by computing a Welch's T-statistic and visualized by recoloring each bin by its associated T-statistic. Redder values indicates faster oscillations in that region of expression space, and bluer values indicate slower oscillations in that region of expression space. Note that the diagonal (self-comparison) is grey as expected and the plots are symmetric across the diagonal (up to a sign flip to the statistic).

(B) Effects of the choice of different distance metrics on the appearance of control, ectoderm, mesoderm, and endoderm differentiation trajectories in multi-dimensional scaling (MDS) space. Pairwise distances between portraits defined by Welch's T statistic, Cohen's D, or absolute change in frequency were computed as:

$$T_{welch} = \frac{\overline{X_1} - \overline{X_2}}{\sqrt{\frac{s_1^2}{n_1} + \frac{s_2^2}{n_2}}} \qquad d_{cohen} = \frac{\overline{X_1} - \overline{X_2}}{s_{pooled}} \qquad \Delta freq = \overline{f_1} - \overline{f_2}$$

squared and summed across all bins, and used as input similarity metrics for MDS rescaling and plotted in 3D, with the x and y-axes corresponding to MDS dimensions and the z-axis corresponding to lab-reference time. All three metrics resulted in similar qualitative trajectories in the Waddington-like MDS space.

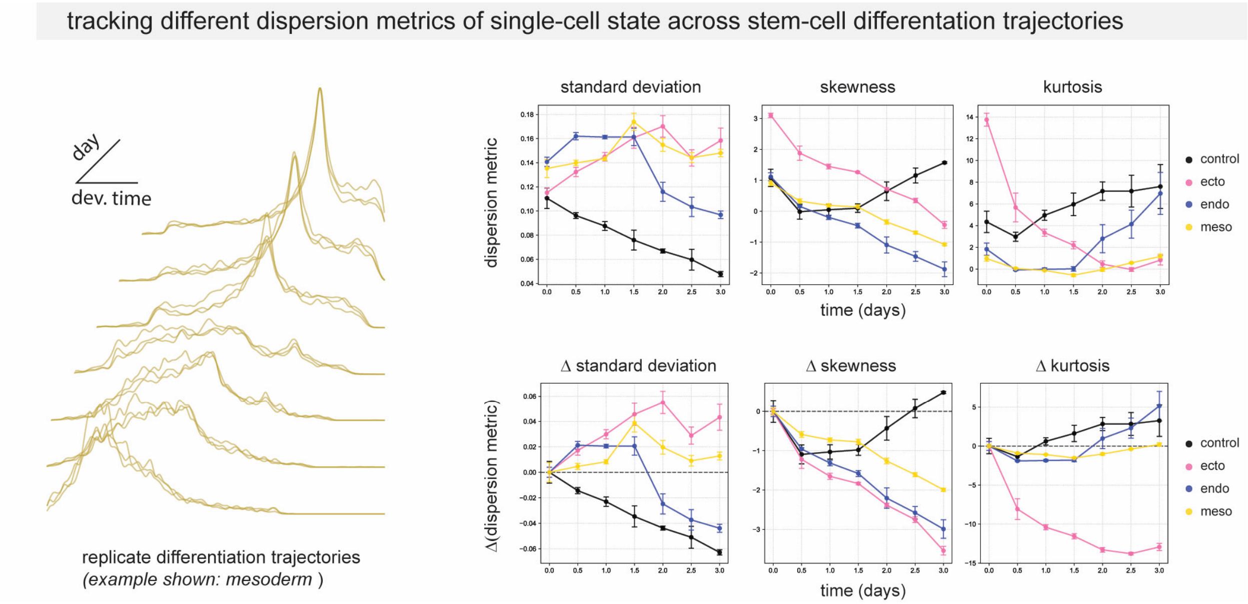


**Figure S12. Dispersity metrics of single-cell heterogeneity during stem-cell differentiation into different cell types.**

Following single-cell developmental progression estimation for all cells across all experiments, the developmental heterogeneity within each population's distribution over time during differentiation was explored by computing the absolute and relative (to the day 0 timepoint) statistical moments of the distribution.

**Supplemental Movie Legends.**

**Movie S1. Genetically encoded oscillators (GEOs) generate synthetic fluorescent carrier signals for single-cell streaming and data encoding.** Representative videos of GEO-expressing cells and cell-populations as well as workflows for frequency domain analysis of underlying GEO FM-barcodes and waveform encoding rules using Fast Fourier Transform and aggregation of data from thousands of individual cells.

**Movie S2. GEOs generate persistent single-cell oscillations with defined frequency and amplitude.** Representative low magnification widefield composite fluorescence image of the *E. coli* GEO is shown. FFT-based image processing transforms time series data into frequency-domain data, producing an image power spectrum that illustrates the distribution of oscillatory power across different frequencies. By false coloring each slice of the multi-channel image power spectrum according to peak oscillatory frequency, a single image is generated to visualize cells with different frequencies within the population. Frequency re-colored images of different species-matched GEO pairs.

**Movie S3. Multi-component GEOs generate a single waveform with uniform frequency.** Representative fluorescence image of a U-2 OS cell transduced with a multi-component GEO system composed of a MinD ATPase (*E. coli*), a slow (*E. coli*) MinE activator, and a fast (*P. damselae*) MinE activator. A representative kymograph, pixel-level time series, and the associated FFT power spectrum for this cell is shown. The image power spectra for this cell at the 77 mHz frequency slice is shown.

**Movie S4. Design and analysis of GEO-FM frequency-modulation single-cell data-encoding circuits.** GEO-FM circuits enable the characterization and quantification of transcription factor activity or protein degradation dynamics at a single-pixel resolution, offering a detailed, localized readout of cell state behavior. Pixel-level data can be aggregated at the cellular level to generate single-cell response trajectories for cell-state data.

**Movie S5. GEO performance in human embryonic stem cells and differentiated lineages.** Representative videos of GEO-expressing human embryonic stem cells (hESCs) and differentiated germ layer populations (three days post differentiation) and associated single-cell barcodes. Frequencies are colored as in the colormap from Figure 5. Videos are shown at 60x speedup. Scale bar: 40 um.

**Movie S6. GEO autoencoder analysis of stem cell differentiation trajectories and single-cell developmental heterogeneity.** Workflow for quantitative analysis of GEO autoencoder signals from the undifferentiated and differentiated hESC populations in Figure 6. Population waveform encoding behaviors collected across different timepoints are used as anchors for single-cell GEO estimation of developmental progress. Single-cell estimates allow investigation of spatiotemporal heterogeneity during the differentiation timecourse and enable single-cell state estimates to be projected directly onto their corresponding FM-barcoded GEO signals in microscopy images.
